# Sequence independent activity of a predicted long disordered segment of the human papillomavirus L2 capsid protein during virus entry

**DOI:** 10.1101/2023.03.21.533711

**Authors:** Changin Oh, Patrick M Buckley, Jeongjoon Choi, Aitor Hierro, Daniel DiMaio

**Affiliations:** Department of Genetics, Yale School of Medicine, PO Box 208005, New Haven, CT 06520-8005, USA; Department of Microbial Pathogenesis, Yale School of Medicine, New Haven, CT; CIC bioGUNE, Bizkaia Technology Park, 48160 Derio, Spain; IKERBASQUE, Basque Foundation for Science, 48009 Bilbao, Spain; Department of Therapeutic Radiology, Yale School of Medicine, PO Box 208040, New Haven, CT 06520-8040, USA; Department of Molecular Biophysics & Biochemistry, Yale University, PO Box 208024, New Haven, CT 06520-8024, USA; Yale Cancer Center, PO Box 208028, New Haven, CT 06520-8028, USA

**Keywords:** papillomavirus, HPV, intrinsically disordered region, IDR, retromer, L2 protein

## Abstract

The papillomavirus L2 capsid protein protrudes through the endosome membrane into the cytoplasm during virus entry to bind cellular factors required for intracellular virus trafficking. Cytoplasmic protrusion of HPV16 L2, virus trafficking, and infectivity are inhibited by large deletions in an ∼110 amino acid segment of L2 that is predicted to be disordered. The activity of these mutants can be restored by inserting protein segments with diverse compositions and chemical properties into this region, including scrambled sequences, a tandem array of a short sequence, and the intrinsically disordered region of a cellular protein. The infectivity of mutants with small in-frame insertions and deletions in this segment directly correlates with the size of the segment. These results indicate that the length of the disordered segment, not its sequence or its composition, determines its activity during virus entry. Sequence independent but length dependent activity has important implications for protein function and evolution.

## Introduction

The amino acid sequence of proteins determines their structure and function. The folded shape of proteins is specified by their amino acid sequence, although molecular chaperones can assist in folding (Anfinsen, 1973) (Horwich & Fenton, 2020). Many proteins contain domains (intrinsically disordered regions, IDRs) that do not adopt stable folded structures in solution, but the activities of IDRs are also dependent on amino acid sequence or composition. For example, IDRs often contain short specific amino acid sequences that serve as binding sites for other proteins or adopt well-folded structures when they associate with binding partners (Babu, 2016). Even for IDRs that appear to function primarily as linkers or spacers, overall amino acid composition or sequence complexity is important, and often specific sequence features such as the number of prolines or glycines, the net charge, or the distribution of charged residues are key (Das & Pappu, 2013; Zarin et al., 2019) (Zarin et al., 2021). In some cases, IDRs can undergo phase separation, the structural basis for which remains controversial but also appears sequence- or composition-dependent (Fawzi et al., 2021; Kato et al., 2022).

Human papillomaviruses (HPVs) are non-enveloped viruses that trigger ∼5% of cancers (de Martel et al., 2017). Each HPV particle contains 360 molecules of the L1 protein, which forms the outer shell of the capsid, and up to 72 molecules of the L2 protein, which is essential for trafficking of the viral DNA genome to the nucleus via the retrograde transport pathway during virus entry (Siddiqa et al., 2018) (Keiffer et al., 2021) (Buck et al., 2008) (Campos, 2017). L2 contains a highly conserved short C-terminal sequence of basic amino acids that functions as a cell-penetrating peptide (CPP) after virus internalization to drive most of the L2 molecule through the endosomal membrane into the cytoplasm (Zhang et al., 2018) (DiGiuseppe et al., 2015). A short sequence in L2 adjacent to the CPP then binds directly to retromer, a cytoplasmic protein sorting complex that delivers the incoming virus into trans-Golgi network (TGN) and the retrograde pathway for trafficking to the nucleus (Lipovsky et al., 2013) (Popa et al., 2015). The central portion of the L2 protein binds other cytoplasmic trafficking factors such as SNX17 and COPI (Bergant Marusic et al., 2012; Harwood et al., 2020; Harwood et al., 2023), and an N-terminal transmembrane domain (TMD) appears to anchor the L2 protein in the endosomal membrane after L2 has protruded into the cytoplasm (Bronnimann et al., 2013). Only small, unassigned densities are attributed to L2 in cryo-EM reconstructions of HPV16 capsids, and circular dichroism analysis of the HPV16 L2 protein purified from bacteria implies a substantial fraction of the protein is disordered (Goetschius et al., 2021) (Breiner et al., 2019). Upon SDS-polyacrylamide gel electrophoresis, L2 migrates aberrantly slowly. The structural basis for the unusual activities of the L2 protein is not known (Campos, 2017).

Here, we report that much of L2 is predicted to be disordered, including a poorly conserved C-terminal stretch of approximately 110 amino acids located between a conserved central structured region containing the COPI and SNX17 binding sites and the retromer binding site/CPP. This segment of L2 is required for membrane protrusion, retromer association, and proper virus trafficking during entry, but these activities are determined by the length of this segment, not by its amino acid sequence or composition.

## Results

### Much of the L2 protein is predicted to be disordered

We used Alphafold2 (AF2) to generate structural models of the L2 protein from HPV16 and other papillomaviruses (Jumper et al., 2021). AF2 predicts that the 473-residue HPV16 L2 protein contains N-terminal α-helices and a central structured region, but most of L2 is disordered (Fig. 1A). The central structured region consists of a prominent three-stranded anti-parallel β-sheet that contains the required COPI binding site and is immediately C-terminal to a region we named the rollercoaster, which consists of short α-helices connected by loops (Fig. 1B). This central structured region is located between an ∼150-residue N-terminal disordered region and an ∼110-residue C-terminal disordered region (designated NDR and CDR, respectively) (Fig. 1C). The required retromer binding site and CPP are immediately downstream of the CDR. This overall architecture was predicted for all L2 proteins examined, including HPV18 (a genital α-HPV closely related to HPV16); HPV5 (a divergent cutaneous β-HPV), bovine papillomavirus type 1 (an animal δ-fibropapillomavirus), and two of the most divergent known papillomaviruses, CmPV1 and CcPV1 (dyozetapapillomaviruses from sea turtles) (Fig. S1A). The predicted fold of the β-sheet is nearly identical in all these L2 proteins (HPV16 and CmPV1 are shown as examples) (Fig. 1B). The RoseTTAFold structure prediction program also predicted that HPV16 L2 contains a central β-sheet flanked by long disordered segments (Fig. S2) (Jones & Cozzetto, 2015) (McGuffin et al., 2000). The DISOPRED3 and flDPnn disorder probability prediction programs (Hu et al., 2021; Jones & Cozzetto, 2015) also predicted that these segments of L2 are disordered in divergent papillomavirus types, and the AF2 pLDDT confidence score of these segments are low, as is commonly seen for IDRs (Fig. 1C and S3) (Ruff & Pappu, 2021). These predicted disordered regions are poorly conserved and have relatively low sequence complexity, as is the case for many IDRs (Fig. 1C and S1B-C) (Romero et al., 2001).

**Figure 1.**
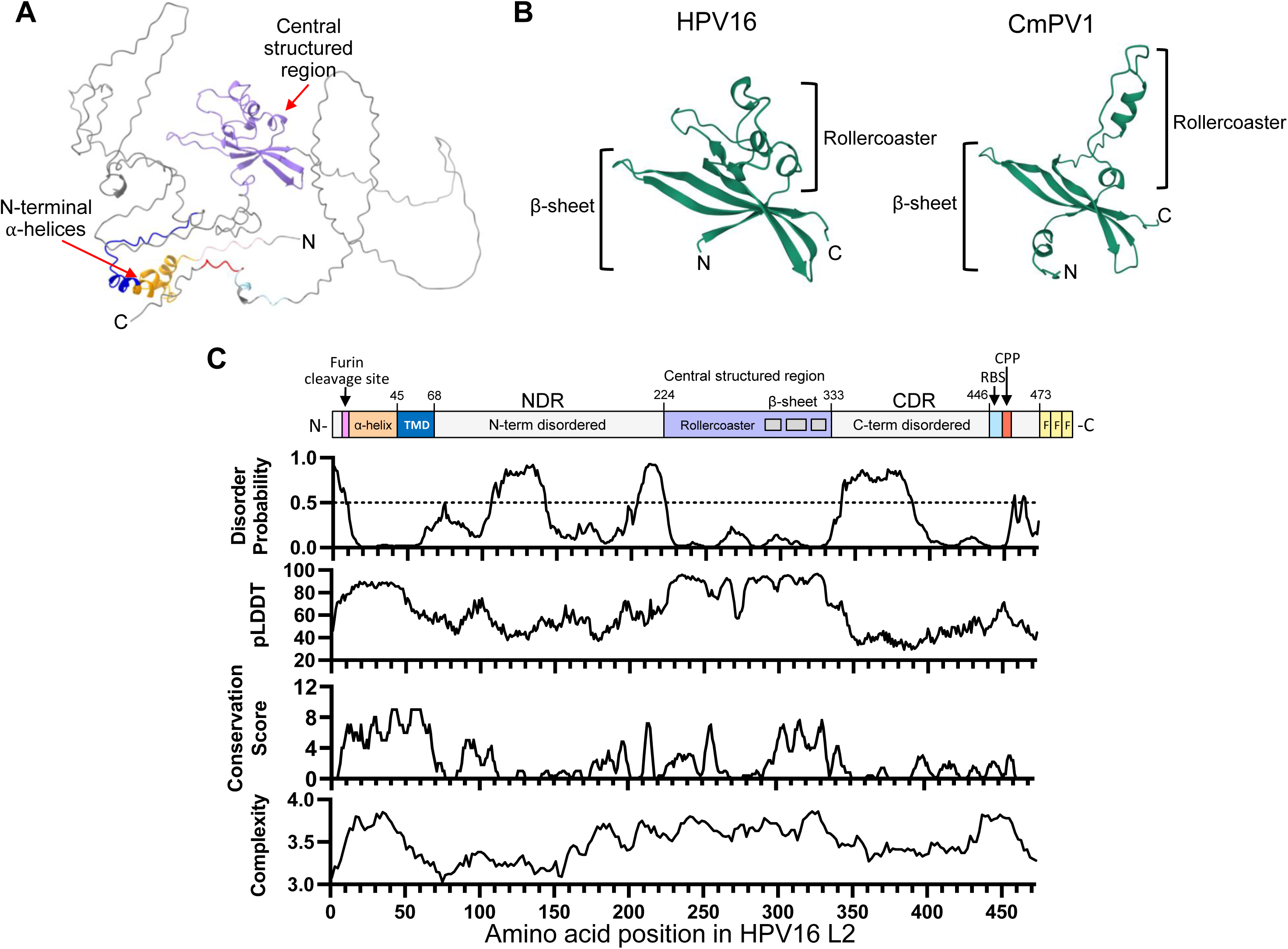
AlphaFold and protein sequence analysis of papillomavirus L2 proteins predicts structured and disordered regions. **A**. AlphaFold2 prediction of HPV16 L2 showing the N-terminal α-helices (orange), transmembrane domain (blue), central structured region (purple) and disordered regions (grey). The N and C termini are indicated. **B.** Comparison of the predicted central structured region of HPV16 and CmPV1 showing the β-sheet and rollercoaster. **C.** Cartoon at top shows a schematic of HPV16 L2 with key amino acid positions numbered. TMD, putative transmembrane domain; NDR, N-terminal disordered region; CDR, C-terminal disordered region; RBS, retromer binding site; CPP, cell penetrating peptide; F, FLAG tag. Graphs are aligned with the L2 schematic above showing disorder probability score from DISOPRED3 (higher values indicate greater likelihood that the segment is disordered), AlphaFold2 pLDDT values (higher values indicate higher confidence in the prediction), five- amino acid rolling conservation score of 18 different HPV types (six each alpha, beta, and gamma) and six animal papillomavirus types calculated by Jalview (higher values indicate more conservation) (viruses included are listed in Figure S1D), and rolling sequence complexity scores (K_2_ values), calculated as described by (Wootton & Federhen, 1996).

### A C-terminal disordered region of L2 is required for infectivity but can tolerate many in-frame mutations

To determine whether the CDR is required for infection, we tested the infectivity of HPV16 pseudoviruses (PsVs) containing L2 with large deletions in this region (Δ337-385 and Δ386-441) (Fig. 2A). PsVs composed of L1 and FLAG-tagged L2 containing a reporter plasmid expressing green fluorescence protein (GFP) were produced in 293TT cells, and infectivity was measured by incubating PsV (normalized for packaged reporter plasmid content) with HeLa S3 cells and measuring GFP fluorescence by flow cytometry at 48 hours post infection (hpi). This assay measures virus entry and trafficking to the nucleus but not later stages of the virus life cycle. Even though the deleted L2 proteins were assembled into capsids (Fig. S4), both deletion mutants were non-infectious, showing that this region of L2 is required for infection (Fig. 2A). Surprisingly, however, infectivity was not inhibited by a nine-amino acid HA tag inserted at various positions in the CDR (Fig. 2B). Similarly, infectivity was not substantially inhibited by four-to fifteen-residue deletions that in aggregate spanned the entire region between 337 and 445 (Fig. 2C and S5A). In contrast, activity is abolished by deletion of residues 330 to 336, which span the boundary between the predicted central structured region and the CDR. Based on these results, we define the CDR as amino acids 337 to 445 of HPV16 L2. This mutational analysis suggests that there are no unique sequences or structures in the CDR that are required for infection. Some of the active mutants contained relatively low amounts of L2 (Fig. S4), implying that the number of L2 molecules in these mutant capsids produced in 293TT cells is not limiting for infection.

**Figure 2.**
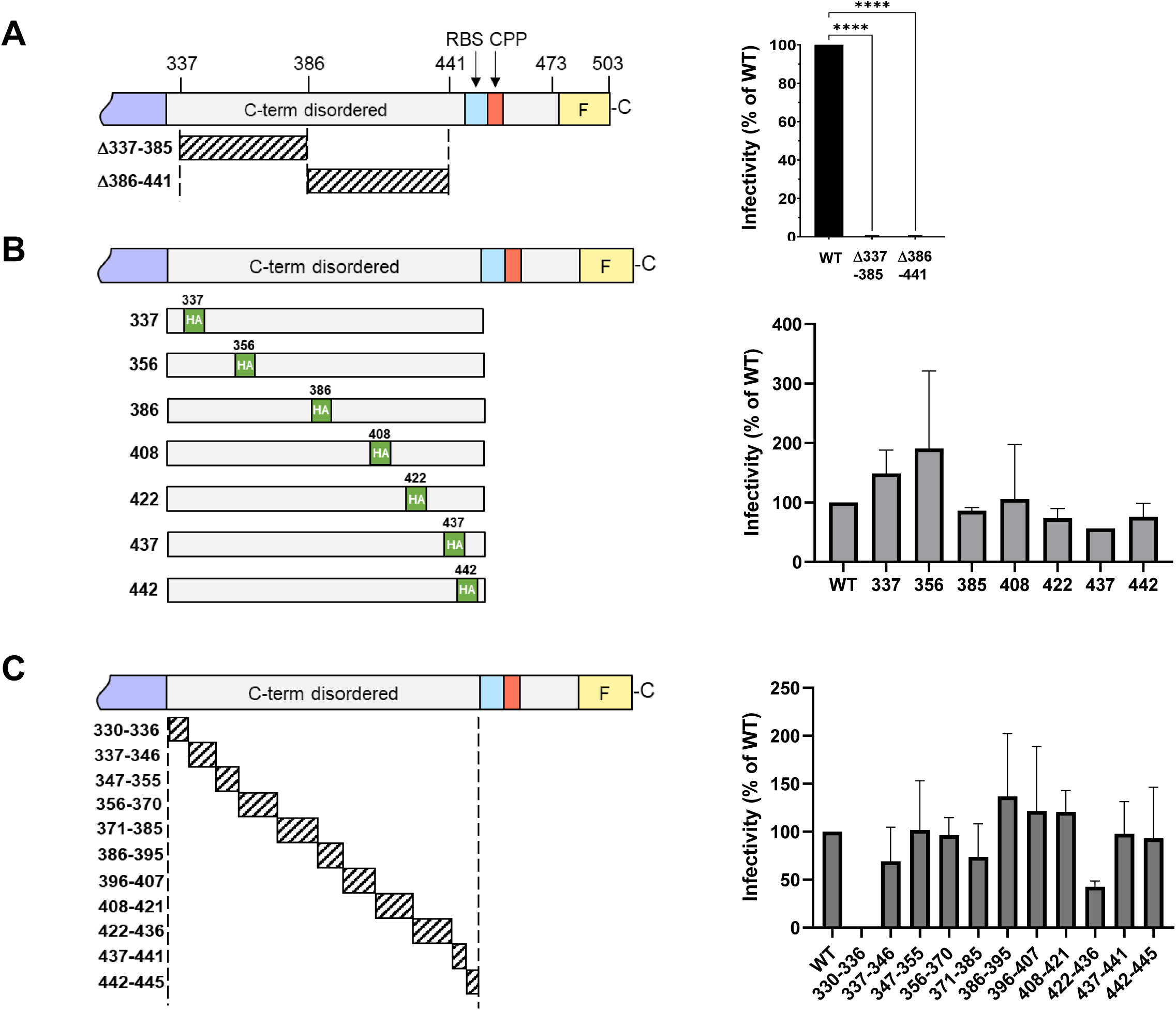
HPV L2 CDR is required for infection but tolerates small in-frame insertion and deletion mutations. **A.** Left. Top shows a schematic diagram of the C-terminus of wild-type HPV16 L2 protein as in Figure 1C, except 3X FLAG tag is represented by a single F. Large in-frame deletions are represented with hatched rectangles. Right. HeLa S3 cells were infected with wild-type (WT) or mutant PsV containing equal numbers of the GFP reporter plasmid (corresponding to MOI of one for wild-type). GFP fluorescence was measured by flow cytometry 48 hours post infection (hpi) and mean fluorescent intensity used to calculate relative infectivity, expressed as percent of wild-type HPV16 PsV (set at 100%). Mean results and standard deviation of three independent experiments are shown. ****, p-value < 0.0001 compared to wild-type as assessed by one-way ANOVA. **B.** As in panel A, except the positions of in-frame HA tag insertions are represented by the green rectangles. None of the mutants displayed statistically different activity from wild-type as assessed by one-way ANOVA . **C.** As in panel A, except small in-frame deletions are represented by the hatched rectangles. None of the mutants between positions 337 and 441 are statistically different from wild-type as assessed by one-way ANOVA. Mutant Δ330-336 was significantly impaired (p-value < 0.0001)

### Insertion of diverse sequences can rescue the activity of L2 mutants with large deletions in the C-terminal disordered segment

We next tested if the infectivity defect caused by the large deletions in the CDR could be rescued by inserting other sequences (Fig. S5B and C). First, we generated 49SR and 56SR by replacing each large deletion with randomly scrambled amino acids while maintaining the wild-type amino acid composition and length (Fig. 3A). Both replacements rescued infectivity, showing that there is no specific required sequence of amino acids in the CDR (Fig. 3B). Papillomavirus CDRs contain a high proportion of serine, threonine, and proline, and unlike most IDRs are enriched in bulky hydrophobic amino acids and depleted in charged residues (Fig. S6) (Romero et al., 2001) (Tompa, 2002). To test whether this amino acid composition was important for activity, we replaced the deleted sequences with tandem arrays of HA tags that restored the wild-type number of amino acids to generate mutants 49HA and 56HA (Fig. 3A). The tag is enriched in aspartic acid, proline, and tyrosine and lacks serine and threonine (Fig. 3C). These segments restored infectivity, even though their composition differs drastically from wild-type (Fig. 3B). These experiments were conducted with PsV containing FLAG-tagged L2. To determine whether the FLAG tag affected these results, we also tested the infectivity of PsV containing untagged wild-type, Δ386-441, or 56HA L2. As was the case for the tagged versions, the untagged version of Δ386-441 was non-infectious and this defect was rescued by the 56-residue HA tag insertion (Fig. S7A). However, replacing the entire disordered region from position 337 to 441 with scrambled amino acids or tandem HA tags caused only a slight increase in activity, which was most evident in the flow cytometry histograms (Fig. S8).

**Figure 3.**
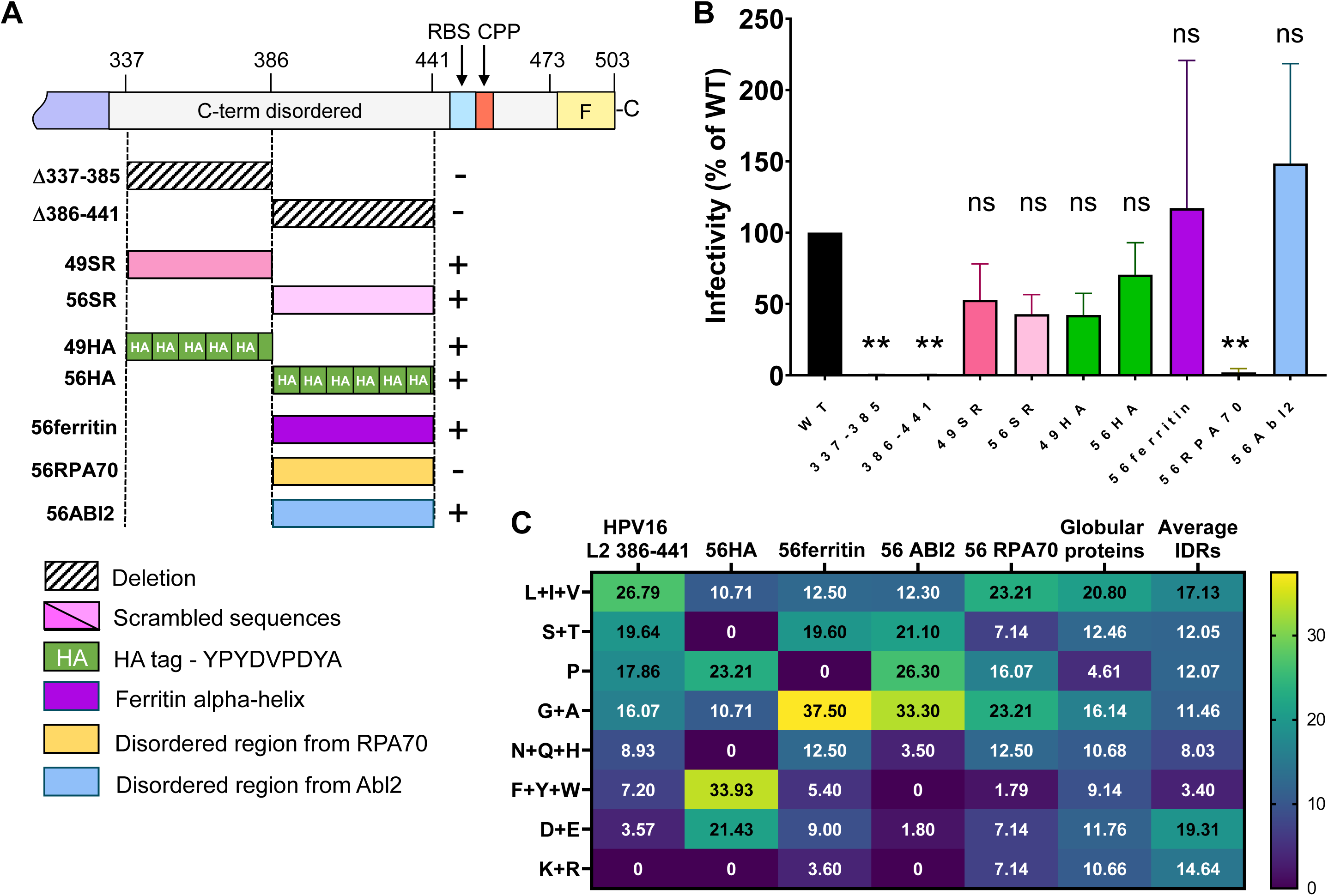
The L2 CDR can be replaced with diverse unrelated sequences. **A.** Top shows schematic diagram of the C-terminus of the wild-type HPV16 L2 protein as in Figure 2A. Below are rectangles representing the large deletion mutations (hatched), scrambled replacements (light and dark pink), tandem HA tag replacements (green), ferritin replacement (purple), RPA70 IDR replacement (yellow), and ABI2 IDR replacement (light blue). The + and – indicate the infectivity of the PsV containing the indicated L2 protein. **B.** Graph of the infectivity of wild-type or mutant PsV was assessed and displayed as in Figure 2. **, p-value< 0.01 compared to wild- type as assessed by one-way ANOVA. ns, not significant. **C.** Heatmap showing the amino acid composition of the C-terminal half of HPV16 L2 CDR (amino acid positions 386-441), replacement sequences, and large compilations of globular proteins and IDRs (Zarin et al., 2019). Each row represents a group of amino acids based on similar chemical characteristics, shown in descending order based on their abundance in the HPV16 L2 segment. Numbers show percent abundance of each group of amino acids in the segment.

For the remainder of this paper, we focused on the largest segment we successfully replaced, the 56-residue segment between position 386 and 441. Because this L2 segment and the HA tags contain numerous proline residues (Fig. 3C and S6), we tested the activity of a proline-free segment derived from short helical segments of ferritin separated by glycine-rich linkers (Fig. 3C). This sequence also rescued infectivity of Δ386-441 (Fig. 3B). Finally, we replaced the segment deleted in Δ386-441 with a 56-residue segment from a known IDR from two human proteins, Abl interactor 2 or Replication Protein A70 to generate 56ABI2 and 56RPA70, respectively (Fig. 3A). Strikingly, the ABI2 segment fully rescued infectivity, demonstrating that L2 activity could be restored with an unrelated sequence that is devoid of fixed structure (Fig. 3B). The IDR from RPA70 did not rescue activity (Fig. 3B), showing that not all IDRs were active. Similar results for all these constructs were obtained in HaCaT human skin keratinocytes (Fig. S7B). The hydrophilicity, isoelectric focusing point, sequence complexity, and net charge of the CDR of representative papillomaviruses and foreign sequences that rescued activity vary widely and do not consistently differ from the RPA70 IDR, which did not rescue (Fig. S9). These results indicate that while some sequences are inactive, several highly diverse sequences from various sources can substitute for naturally occurring viral sequences to support HPV entry.

### The length of the C-terminal disordered region controls infectivity

To determine whether the length of the CDR was important for HPV infection, we tested the infectivity of five nested deletion mutants containing a fixed C-terminus at position 441 (Fig. 4A). As shown in Fig. 4B, there is a direct relationship between the length of this region and infectivity. We next inserted different numbers of HA tags into Δ386-441 (Fig 4A). Again, infectivity was directly proportional to the number of inserted HA tags (Fig. 4C). The activity of these mutants was remarkably congruent despite the markedly different amino acid sequences and compositions of the insertions and deletions (Fig 4D). We conclude that the length of the L2 CDR controls HPV infectivity.

**Figure 4.**
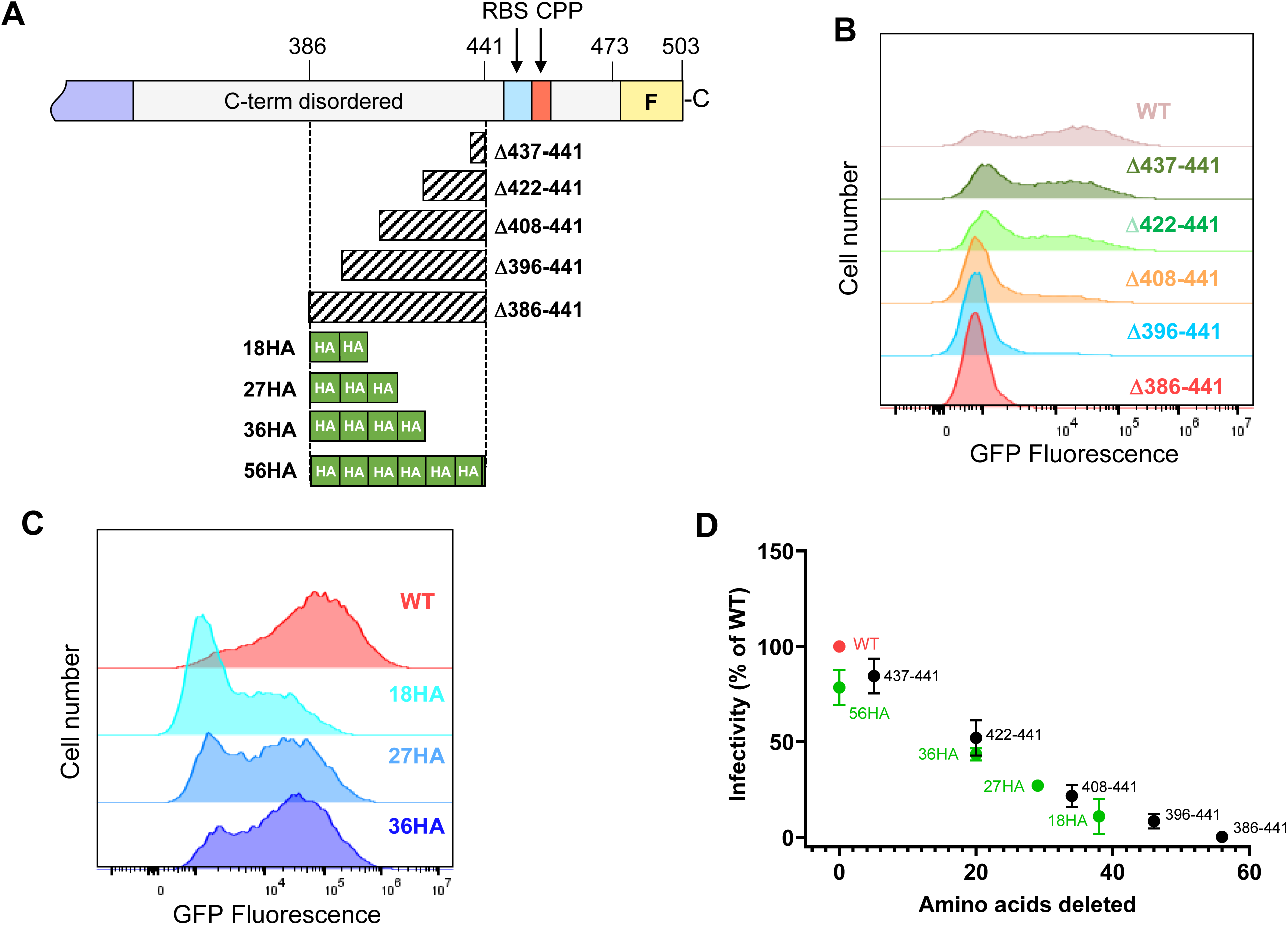
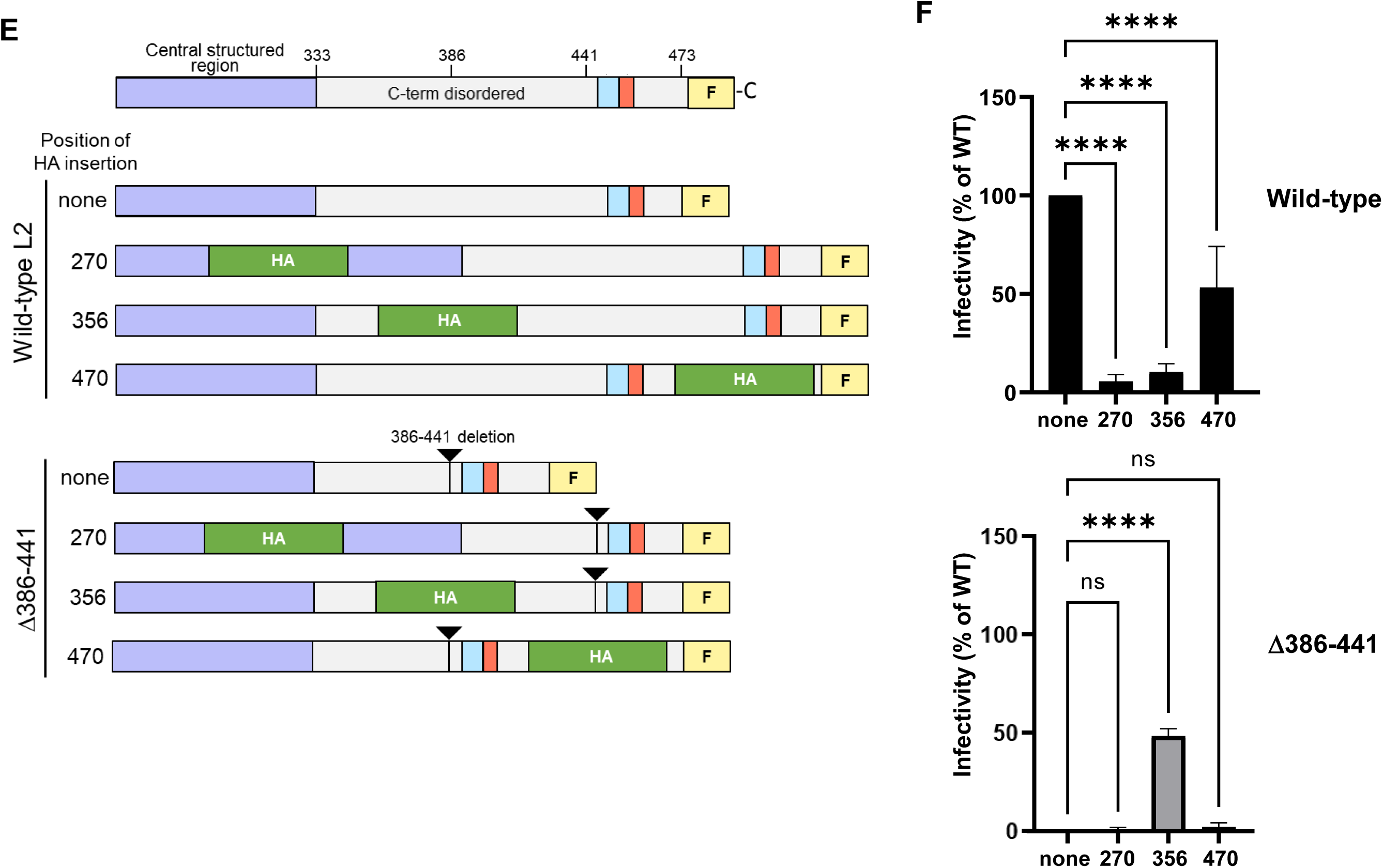
Length of the L2 CDR determines activity. **A.** Schematic diagram of the C-terminus the HPV16 L2 protein with nested deletion (hatched) and HA tag insertion (green) mutations represented below. **B.** Flow cytometry histograms of GFP fluorescence of HeLa S3 cells infected as in Figure 2 with wild-type and nested deletion mutant PsVs, measured at 48 hpi. Similar results were obtained in three independent experiments. **C.** As in panel B but for HA tag insertion mutants. Similar results were obtained in three independent experiments. **D.** Scatter plot of the infectivity of wild-type (red), nested deletion (black) and nested HA insertion (green) mutants versus the number of amino acids deleted compared to wild-type. Each dot represents mean value and standard deviation for each PsV from three independent experiments. **E.** Schematic diagrams of the C-terminal half of wild-type and mutant HPV16 L2. The mutants contain a 56-residue insertion of a tandem array of HA tags (green) in the context of wild-type or Δ386-441 L2, as indicated. The position of the deletion is indicated by the black triangles. **F.** HeLa S3 cells were infected at a MOI of 10 (for wild-type), and infectivity was assessed by flow cytometry and displayed as in Figure 2A. The position of the HA insertion is noted at the bottom, as is whether the insertion was in the wild-type or Δ386-441 context. ****, p-value< 0.0001 compared to cells PsV lacking an HA insertion as assessed by one-way ANOVA. ns, not significant.

To determine whether the 56-residue HA insertion rescued infectivity of Δ386-441 by restoring the length of the CDR itself or by restoring the overall length of the L2 protein, we inserted the 56-amino acid HA tag array at the three different positions in wild-type L2: at position 270 in the rollercoaster, at position 356 within the CDR, and at position 470 downstream of the CPP (Fig. 4E). The mutant L2 proteins were assembled into the purified PsV stocks (Fig. S10A). As shown in Figure 4F, the insertion was well-tolerated at position 470. At positions 270 and 356, the insertion caused ∼10-15-fold infection defect, but infectivity was readily detected if infection was performed at high multiplicity of infection (e.g., see Fig. S10B). We also inserted the HA tag array at the same positions in the Δ386-441 context. The HA insertion at position 356 in the CDR largely rescued the infectivity of Δ386-441, but HA inserted at position 270 or 470 did not rescue (Fig. 4F and S10B). These results show that the length of the segment of L2 between the central structure and retromer binding site controls HPV infectivity. We note that the mutant containing both Δ386-441 and the HA insertion was more infectious than the insertion mutant alone (Fig. 4F), possibly because there is an upper size limit to the length of the CDR.

### Deletions in the C-terminal disordered region disrupt retromer association and proper virus trafficking during entry

To test whether a large deletion in the CDR caused a trafficking defect during virus entry, we used proximity ligation assay (PLA) to examine the fate of the Δ386-441 mutant and the rescued 56SR variant in infected HeLa S3 cells. In PLA, a fluorescent signal is generated when HPV is proximal to a marker for a cellular compartment (Lipovsky et al., 2013). The PLA signal for L1 and the TGN marker TGN46 at 20 hpi demonstrated the presence of wild-type and 56SR HPV16 PsV at the TGN, whereas minimal signal was detected in cells infected with Δ386-441 (Fig. 5A), suggesting that the deletion caused a trafficking defect during HPV entry. PLA for L1 and the endosome marker EEA1 at 20 hpi showed that wild-type and 56SR PsVs were mostly absent from the endosome, consistent with their departure to the TGN, whereas Δ386-441 accumulated in the endosome, showing that this mutant was endocytosed but impaired for endosome exit (Fig. 5B).

**Figure 5.**
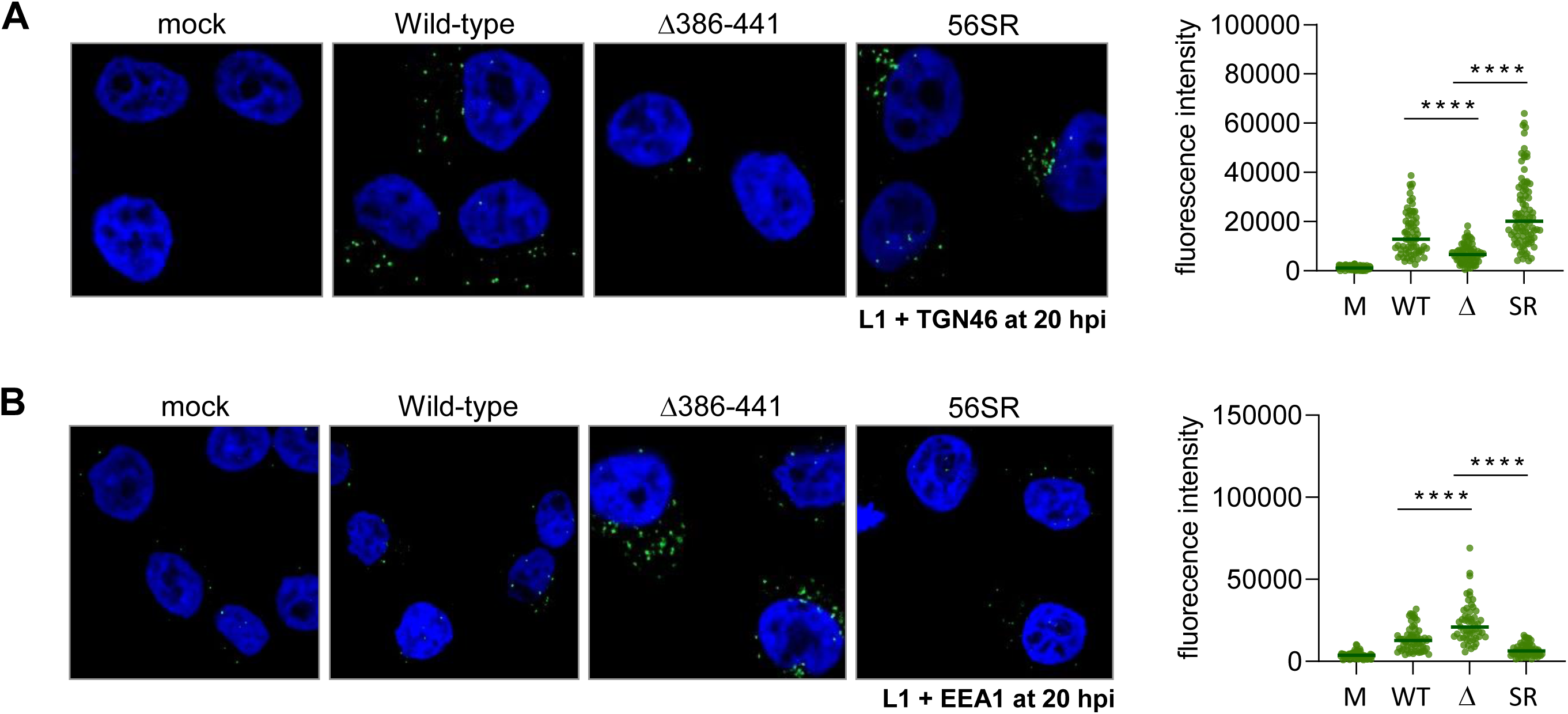
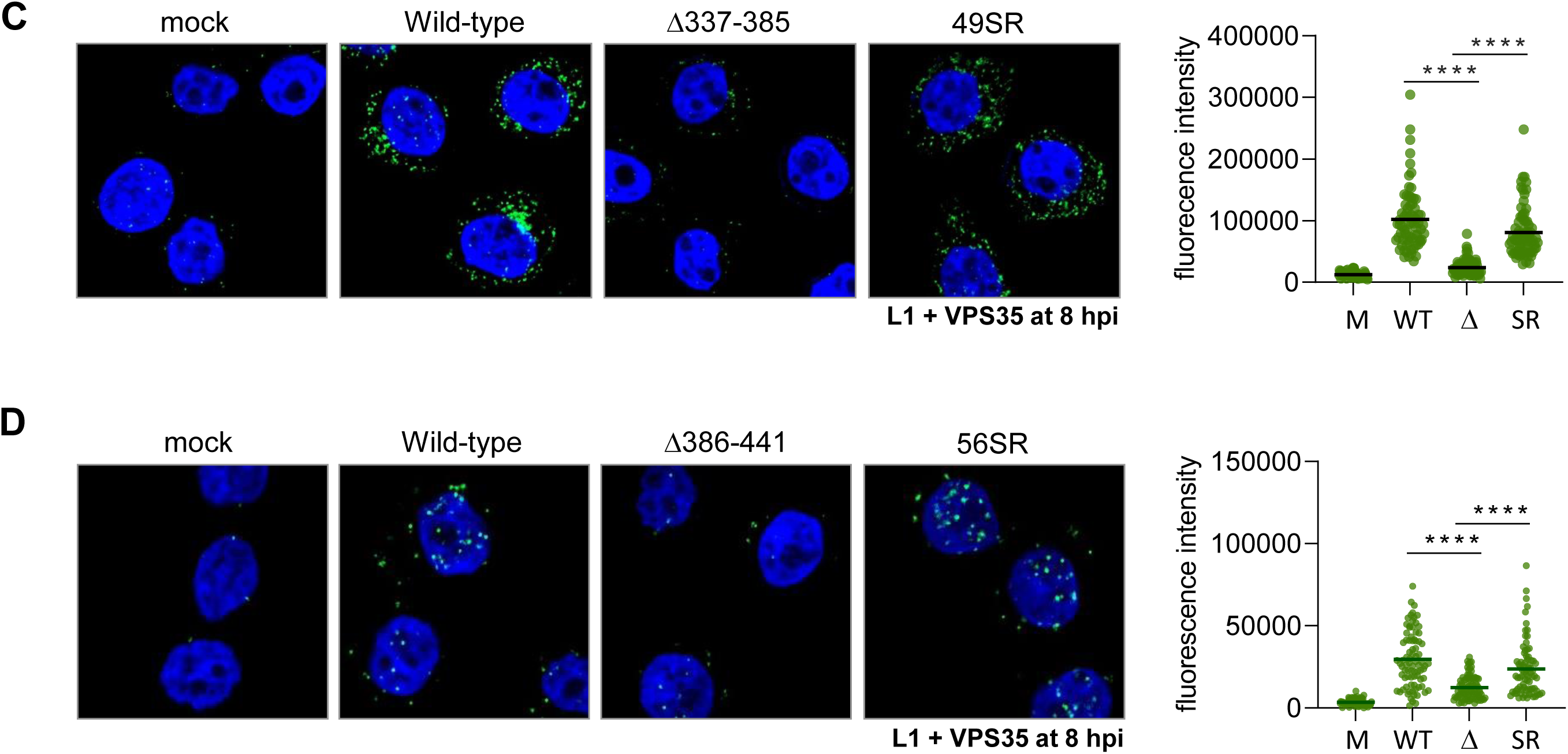
Large deletion in the CDR inhibits association with retromer and endosome exit. **A.** Left, Hela S3 cells were mock-infected or infected at MOI of 200 with HPV16 PsV containing FLAG-tagged wild-type, Δ386-441, or 56SR L2. At 20 hpi, PLA was performed with antibodies recognizing HPV16 L1 and trans-Golgi marker TGN46. Nuclei were stained with DAPI (blue). PLA signal (green) from at least 50 cells for each condition was quantified by ImageJ software. There was no PLA signal in uninfected cells. Right, the quantified PLA result was plotted using GraphPad Prism 9. Each dot represents the PLA fluorescence intensity of an individual cell. Statistical significance was assessed by ANOVA. ****, p-value < 0.0001. Similar results were obtained in three independent experiments. **B.** As in panel A, except PLA was performed with antibodies recognizing L1 and endosome marker EEA1. **C.** As in panel A, except cells were infected with HPV16 PsV containing FLAG-tagged wild-type, Δ337-385, or 49SR L2. PLA was performed at 8 hpi with antibodies recognizing L1 and retromer subunit VPS35. **D.** As in panel C except cells were infected with wild-type, Δ386-441, and 56SR PsV.

Because an endosome accumulation defect is caused by a retromer binding site mutation in L2 (Popa et al., 2015), we performed PLA for L1 and retromer subunit VPS35 to test whether Δ337-385 and Δ386-441 displayed impaired retromer association (Fig. 5C and 5D). As expected, the L1-VPS35 PLA signal at 8 hpi showed that wild-type, 49SR, and 56R PsVs were in close proximity to retromer at this time. In contrast, Δ337-385 and Δ386-441 showed significantly reduced retromer association. These findings show that large deletions in the CDR of L2 interfere with retromer binding and virus trafficking and that these defects are rescued by replacing the wild-type sequence with the same number of scrambled amino acids.

The retromer binding site nominally extends from L2 positions 446 to 455 and is thus retained in the Δ386-441 mutant (Fig. S5A and S11A). To confirm that the retromer association defect of Δ386-441 during infection was not due to the inability of the remaining L2 sequences to bind retromer, we tested whether an L2 peptide lacking sequences N-terminal to position 442 could bind retromer *in vitro*. We incubated detergent extracts from uninfected HeLa cells with biotinylated peptides (L2 amino acids 442-461) that contain a wild-type or a mutant retromer binding site (Fig. S11A). After pulling down the peptide and associated proteins with streptavidin, and we performed western blotting with an antibody recognizing the VPS26 subunit of endogenous retromer. As shown in Fig. S11B, as expected, no VPS26 precipitated in the absence of peptide. VPS26 was pulled down with the peptide containing the wild-type retromer binding site but not by the peptide with the mutant site. Thus, in this peptide binding assay, sequences absent from the Δ386-441 deletion mutant were not required for retromer binding, showing that the deletion that inhibited retromer association in infected cells did not directly prevent retromer binding.

### Large deletions in the C-terminal disordered region inhibit cytoplasmic protrusion of L2

Because protrusion of L2 through the endosomal membrane during infection is required for retromer binding, we tested whether a large CDR deletion inhibited L2 protrusion. Using an established immunofluorescence protocol (DiGiuseppe et al., 2016), we infected HeLa cells with various PsVs, and at 8 hpi we treated cells with saponin, which permeabilizes all cell membranes, or with a low concentration of digitonin, which disrupts the plasma membrane without allowing antibody access to the late endosome lumen. We then performed immunofluorescence staining for L1 and for the FLAG tag on the C-terminus of L2 and examined the cells by confocal microscopy. As expected, following saponin treatment we detected abundant L1 and FLAG staining in cells infected with each of the PsVs (wild-type, Δ386-441, or the corresponding scrambled replacement mutant, 56SR), whereas in all cases there was minimal L1 staining in digitonin-permeabilized cells because L1 is confined to the endosome lumen (Fig 6). Following permeabilization with digitonin, FLAG staining was readily detected in cells infected with wild-type PsV, indicating that the C-terminus of L2 in the cytoplasm was accessible to antibody, as reported previously (Xie et al., 2020). In contrast, staining was significantly lower in cells infected with Δ386-441 but was restored by replacing the deleted sequences with scrambled amino acids in the active 56SR PsV. Taken together, these results indicate that the large deletion inhibited L2 protrusion and that protrusion activity is determined by the length of the CDR, not its sequence. We conclude that the reduced association of Δ386-441 with retromer and defective trafficking is due to impaired protrusion of the deleted L2 protein through the endosome membrane.

**Figure 6.**
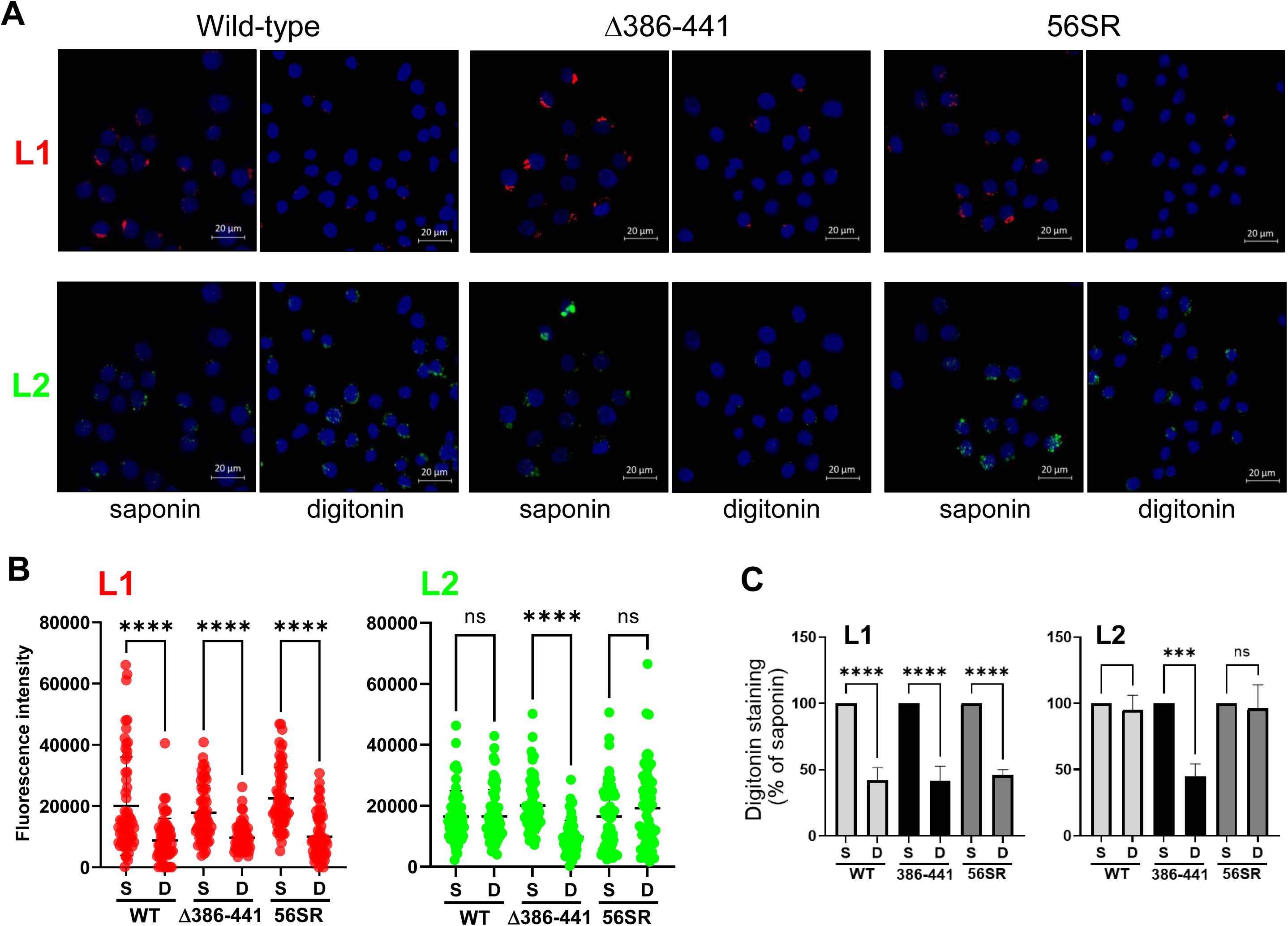
Large deletions in the CDR inhibit protrusion of L2 into the cytoplasm. **A.** HeLa S3 cells were mock-infected or infected at MOI of 100 with HPV16 PsV containing FLAG-tagged wild-type, Δ386-441, or 56SR L2. At 8 hpi cells were permeabilized with 1% saponin or 2 μg/mL digitonin as indicated and immunostained with anti-L1 antibody (red, top row of images) and anti-FLAG antibody (green, bottom row of images). Nuclei were stained with DAPI (blue). There was negligible staining in mock-infected cells. **B.** L1 and FLAG signals as in panel A were quantified for at least 60 cells per condition by using ImageJ. Each dot in the graphs shows the fluorescence intensity of an individual cell. Statistical significance was assessed by using GraphPad Prism 9 to make multiple comparison of each group with one-way ANOVA. ***, p-value <0.001; ****, p-value < 0.0001; ns, not significant. Similar results were obtained in three independent experiments. In addition, in single experiment, protrusion of Δ338-385 PsV was also blocked and restored in its replacement mutant, 49SR (data not shown). **C.** Staining results from three experiments as in panel B are displayed as the average of each experiment plus and minus the standard deviation, expressed as the percentage of saponin staining for each condition.

### Modeling L2 bound to retromer at the endosome membrane

Given the similarity between the FYL retromer binding site in L2 (Popa et al., 2015) and the YLL motif in DMT1-II that bids to the interface between SNX3 and the VPS26 subunit of retromer (Lucas et al., 2016; Tabuchi et al., 2010), we used AlphaFold-Multimer (Evans et al., 2022) to predict whether a peptide that contains the HPV16 L2 retromer binding site (aa 444-GDFYLHPSYYMLR-456) displayed similar binding to retromer as did DMT1-II. The predicted complex positioned the L2 peptide bound to the VPS26-SNX3 interface in a similar position and orientation as in the SNX3–retromer–DMT1-II complex (Fig. 7 and (Lucas et al., 2016)).

**Figure 7.**
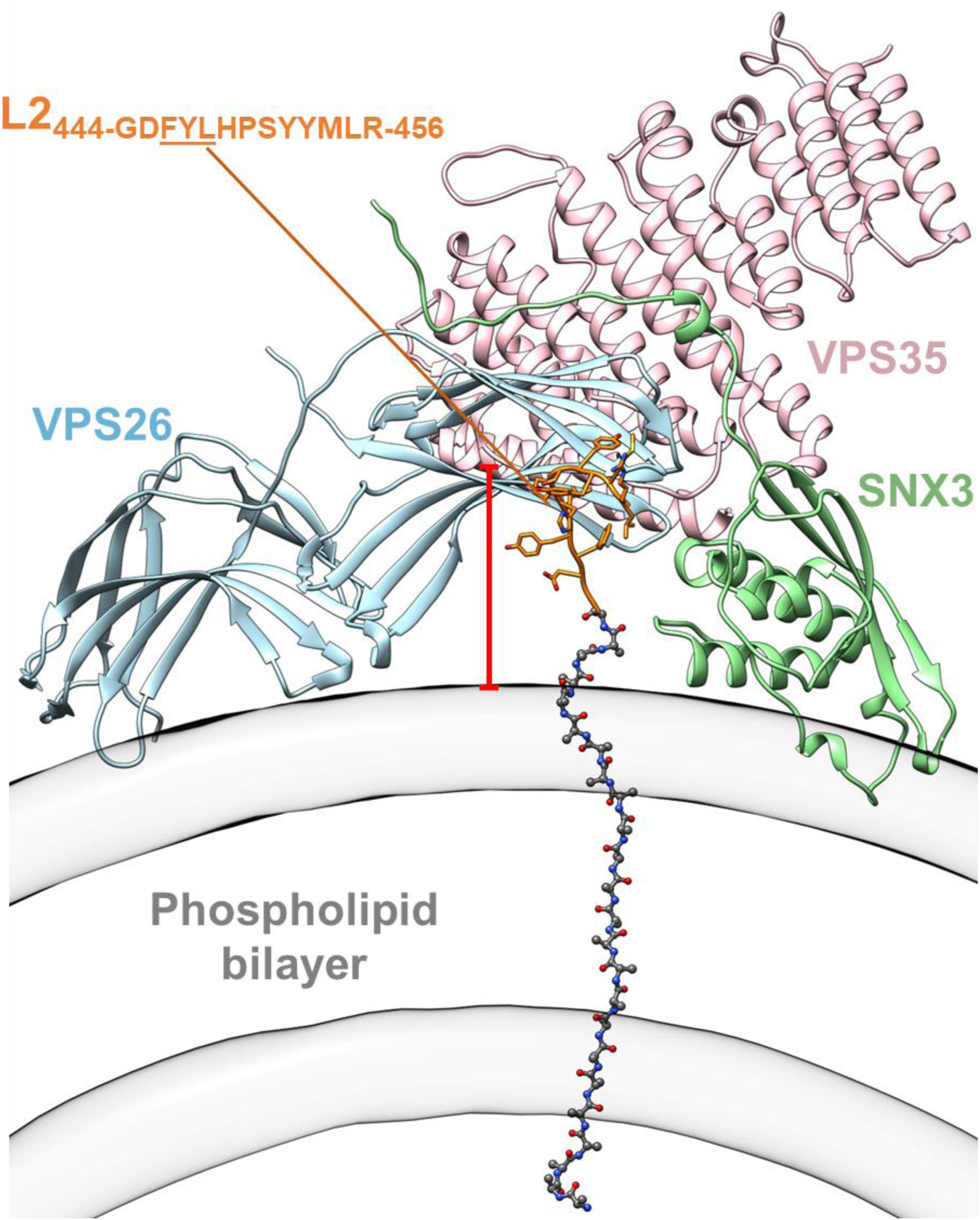
Model of L2 bound to retromer at the endosome membrane. Model predicted by AlphaFold-Multimer (Evans et al., 2022) of the VPS26:VPS35:SNX3 complex bound to amino acids 444-456 of HPV16 L2 (sequence shown in gold). The inner and outer leaflets of the membrane are shown as cylindrical masks derived from the cryo-ET map of the retromer:SNX3 membrane coat (Leneva et al., 2021). The distance from the cargo-binding pocket in VPS26 (Lucas et al., 2016) to the membrane plane is ≈25Å. This distance could be roughly covered by a polypeptide chain of ≈ 9-10 residues in an extended configuration (shown here with a small twist to align it perpendicular to the membrane surface) (Choi et al., 2013). To illustrate the short distance between the membrane and the cargo binding site on VPS26, we added a poly-Ala sequence of 23 residues (in grey) reaching through the membrane to the retromer binding peptide on L2. The red space bar shows 25Å. Thickness of the membrane is ≈ 48Å.

Furthermore, based on the membrane-bound retromer-SNX3 structure (Leneva et al., 2021), the distance from the L2 retromer binding site to the membrane is approximately 25Å, which requires a minimum of 9-10 amino acids to bridge this distance. Considering the cryo-ET cross-section of the membrane (Leneva et al., 2021) and to provide scale, we added an extended poly-alanine stretch across the membrane to the beginning of the L2 peptide used in the modeling (we used poly-alanine because we do not know the true extent or configuration of L2 as it crosses the membrane). Importantly, fewer than 25 amino acids are required to span the distance from the endosome lumen to the retromer binding site. Thus, a 49- or 56-amino acid deletion in the CDR is long enough to prevent the retromer binding site of L2 from protruding into the cytoplasm.

## Discussion

Modeling and amino acid sequence analysis indicate that the L2 capsid protein from all papillomaviruses examined contains a prominent β-sheet in a central structured element flanked by two long disordered regions. These disordered regions presumably account for the aberrant mobility of L2 upon SDS-polyacrylamide gel electrophoresis, because long IDRs often migrate slowly, e.g., (Hanoulle et al., 2009). The C-terminal disordered region of L2 (the CDR) lies between the required COPI binding site in the β-sheet and the retromer binding site and CPP near the C terminus. The CDR is highly tolerant of small in-frame insertion and deletion mutations, but large deletions in the CDR block entry. Importantly, the activity of these large deletion mutants can be rescued by diverse protein segments that differ markedly in amino acid sequence or composition, or in the chemical properties of their constituent amino acids. The activity of mutants containing foreign sequences in either half of the CDR is not due to the presence of redundant sequence elements in the remaining half of the CDR because deleting either half abrogates activity and the defects caused by either deletion can be rescued independently with a different scrambled sequence. Taken together, these findings demonstrate that the PsV entry activity of the CDR is largely sequence independent. Rather, the length of the disordered segment of the L2 protein between the central structured region and the retromer binding site controls infectivity. These conclusions are consistent with the observation that the length of the CDR in different papillomaviruses varies over a relatively narrow range, from about 95 to 130 amino acids, despite poor sequence conservation in this region (Fig. 1C and S1B and S1C). The ability of an IDR to sample an ensemble of conformations ranging from compact to extended may explain why the infectivity of the nested deletion mutants does not display an abrupt length cut-off.

Our results indicate that there are no strict sequence constraints in the CDR for assembly of L2 into PsV, DNA packaging, or virus entry and trafficking to the nucleus. There are reports that this region of L2 plays a role in various activities, including the interaction with L1 (reviewed in (Chen et al., 2023) (Campos, 2017). In fact, a recent paper reports that replacement of three acidic residues at positions 337, 338, and 340 in HPV16 L2 prevents assembly of L2 into the capsid and blocks infection (Chen et al., 2023). We show here that the Δ337-346 mutant and the 49HA and 49SR replacement mutant L2 proteins lack these acidic residues but are assembled into infectious PsV (Figs. 2, 3, and S4). We assume that these differences reflect the particular mutations studied. Most notably, the requirement that the CDR be a certain minimal length will confound studies that delete or truncate the CDR. Regardless, our results show that HPV PsVs that lack substantial native CDR sequences can assemble, traffic to the nucleus, and express an encapsidated genome.

Although either half of the CDR can be functionally replaced with diverse sequences, replacing the entire region with a scrambled or a foreign sequence caused only a modest increase in activity compared to the 105-residue deletion. The sequences that poorly rescue the entire CDR are active when they replace the N-terminal or C-terminal half of the CDR separately, suggesting that these sequences are not intrinsically unable to rescue activity. Further experiments are required to determine why some replacement mutants are not active.

The CDR appears to belong in the entropic chains class of IDRs, which typically display poor amino acid sequence conservation and act as flexible spacers or linkers without adopting a specific structure (van der Lee et al., 2014). However, unlike L2, the specific sequence characteristics or composition of other entropic chains is important for activity. For example, the IDR of RPA70 has a dynamic backbone that is dependent on the amino acid composition and the positions of key glycine residues (Daughdrill et al., 2007). Similarly, the variably sized projection domain of microtubule-associated protein 2 (MAP2) depends on acidic amino acids to generate long-range repulsive forces to ensure the proper spacing of microtubules, and the PEVK domains of the muscle protein titin consists of repeated elements rich in proline, glutamic acid, valine and lysine residues that generate force upon overstretching (Duan et al., 2006) (Mukhopadhyay & Hoh, 2001).

The L2 CDR can be compared to an IDR linker that sets the proper distance between protein binding sites in adenovirus E1A proteins (Gonzalez-Foutel et al., 2022). Both the sequence and the length of this E1A segment are highly variable among different adenovirus strains, but its binding properties and its end-to-end functional length as determined by biophysical methods and modeling are conserved in different adenovirus strains, indicating that during virus evolution, changes in amino acid composition and patterning compensate for differences in sequence length to optimize binding to protein partners. In the case of L2, several unrelated sequences that did not evolve to support virus entry can replace large segments of the CDR, the length of the CDR is not highly variable among different papillomaviruses, and the number of amino acids in CDR determines activity irrespective of sequence. However, the amino acids at certain positions can be aligned in CDRs from different papillomaviruses, and HPV types fall into two clusters, α-types with relatively short CDRs and NDRs and β- and γ-types with relatively long CDRs and NDRs (Fig. S1C and S12). These observations imply that, despite its ability to tolerate foreign sequences, the sequence of the CDR has been subjected to selective pressure during virus evolution, perhaps for L2 activities that are not captured by the assays used here.

During papillomavirus entry, the L2 protein protrudes through the endosomal membrane into the cytoplasm. This unusual feature is key to understanding the role of the CDR in infection and in interpreting the dependence of the CDR on length rather than sequence. The presence of a disordered region N-terminal to the CPP may facilitate the passage of L2 through the endosomal membrane much like the absence of knots permits the passage of a shoelace though an eyelet. Similarly, an IDR adjacent to the retromer binding site may facilitate retromer binding, given that retromer binding sites in cellular cargos often are present in disordered regions (Mukadam & Seaman, 2015).

Large CDR deletions impair the protrusion of the truncated L2 protein into the cytoplasm of infected cells and inhibit the ability of the incoming PsV to come into proximity to retromer. Because the CDR is immediately N-terminal to the retromer binding site, much of the CDR is likely to be within the membrane or the endosome lumen when L2 first protrudes through the membrane and retromer binding occurs (Fig. 7). We conclude that L2 proteins with large deletions in the CDR are too short to protrude far enough into the cytoplasm with sufficient flexibility to bind retromer. The impaired protrusion of the deletion mutant implies that L2 protrusion proceeds sequentially from the C-terminus, with the CPP being exposed first, followed by the retromer binding site and then the CDR. Thus, the spacing of specific sequence elements in L2 dictates when they are exposed in the cytoplasm during protrusion, ensuring that cellular proteins bind to L2 in the proper order and with the proper timing to act during entry. This arrangement allows retromer to bind near the C-terminus of L2 immediately upon protrusion of the CPP to mediate endosome-to-TGN transport. Protrusion of the entire CDR through the membrane must then occur before the central region of L2 is exposed and binds COPI, which mediates later steps of entry (Harwood et al., 2023).

Most of the L2 protein including sequences N-terminal to the CDR is thought to protrude into the cytoplasm during normal infection (DiGiuseppe et al., 2015), but our finding that the large CDR deletion prevents protrusion implies that N-terminal segments of L2 do not protrude if the CDR is too short. We speculate that portions of L2 remain tethered to the membrane or to the rest of the capsid or some other luminal structure until successful protrusion of the C-terminus releases these constraints or that successful protrusion of the C-terminus actively promotes protrusion of N-terminal segments of L2. Consistent with this conclusion, we previously reported that retromer binding stabilizes membrane insertion and cytoplasmic protrusion of L2 (Xie et al., 2021).

What sort of protein-mediated activities can occur with few if any amino acid sequence constraints? Protein segments that function in a sequence independent manner obviously lack any required structural elements and most likely are disordered. They can act as linkers or spacers that connect or separate elements that flank the disordered region to ensure the proper spacing of these elements for optimal binding to other proteins or for proper folding of the elements themselves. These protein segments could also act as timers during directional processes (such as protrusion through a membrane or emergence from the ribosome during translation), where too short or too long a segment will accelerate or delay a process with deleterious effects. Sequence independent protein segments could also serve a measuring function, where a certain length is required to gain access to a binding partner or execute a biochemical function, or it may impart sufficient conformational flexibility for activity.

The existence of sequence independent protein segments has interesting evolutionary implications. Many protein segments that initially displayed sequence independent activity may have been fine-tuned or optimized during evolution, leading to the sequence and compositional biases present in known entropic chains. In the absence of selection for amino acid sequence or composition, protein segments that act in a sequence independent fashion will be difficult to identify by *in silico* analysis for recurring amino acid motifs or patterns, although they may be identifiable because they accumulate synonymous and non-synonymous mutations at similar rates. Importantly, protein segments displaying sequence independent activity are relatively free to evolve unrelated, additional sequence dependent activities so long as the evolved sequences retain the proper length and do not disrupt the sequence independent activity. If these newly evolved activities arise in alternative translational open reading frames, overlapping genes would result. Such events may be particularly important for viruses, which need to maximize the amount of information encoded by their small genomes.

Finally, there are practical implications of the ability of the CDR to tolerate many different sequences. It will likely be possible to replace portions of the CDR with a variety of sequences to confer novel properties to the virus or the L2 protein without blocking its ability to support infection. For example, it may be possible to insert sequence elements into L2 to probe aspects of L2 function or HPV infection or to deliver short protein binding sites or immunogenic epitopes into the cytoplasm.

## Acknowledgments

Flow Cytometry was conducted in the flow cytometry shared resource of the Yale Cancer Center. We thank Nicolas Fawzi and Stevan Hubbard for helpful discussions, Bayan Galal for assistance, and Nosha Vega for help preparing this manuscript. PMB was supported by a T32 training grant (T32AI055403) and a National Science Foundation Predoctoral Fellowship (DGE- 2139841). This work was supported by a grant from The Spanish Ministry of Science and Innovation to AH (PID2020-119132GB-I00) and a grant from the National Institutes of Health to DD (R35-CA242462).

## Supplemental Figure legends

**Figure S1.**
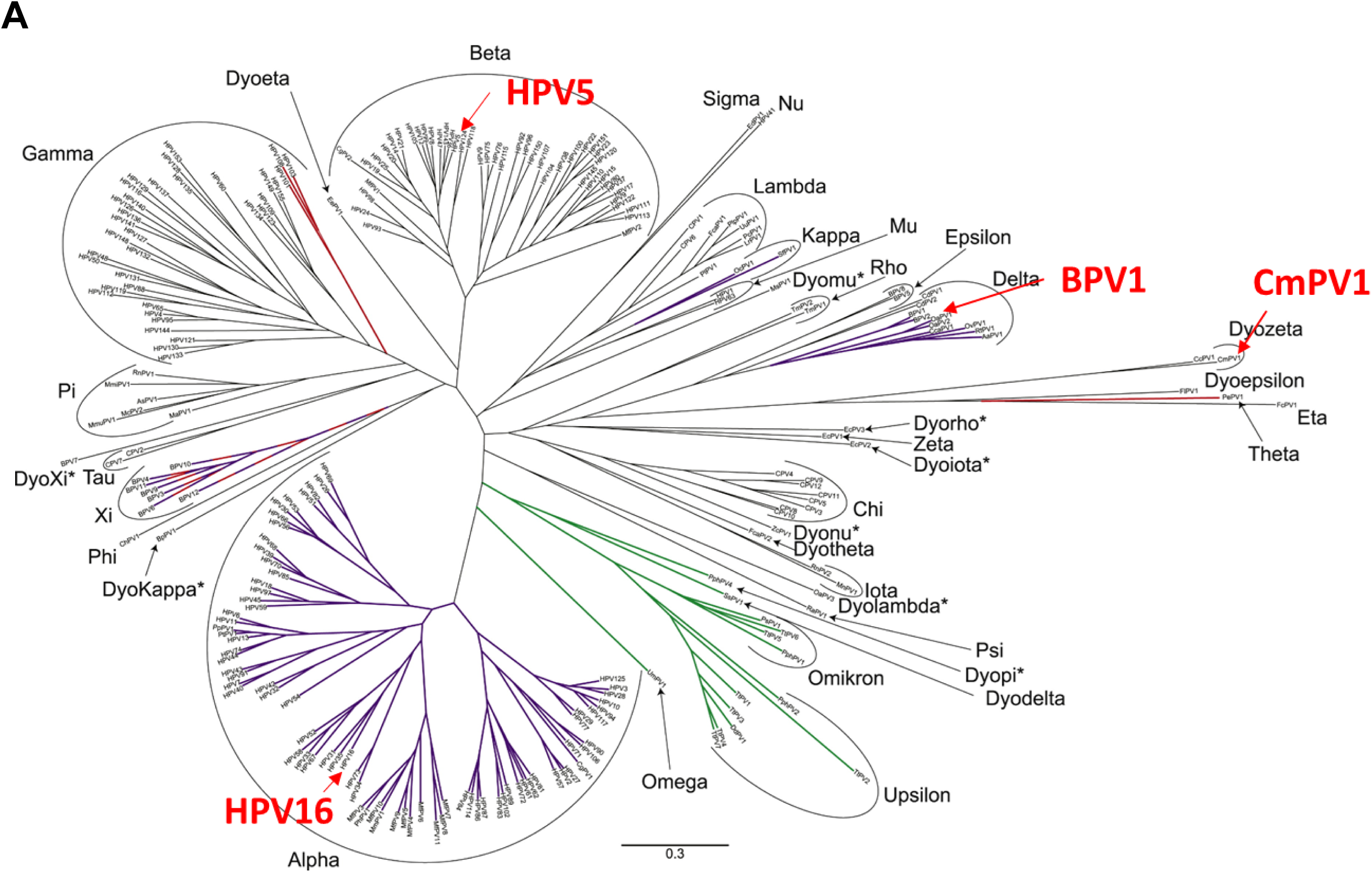
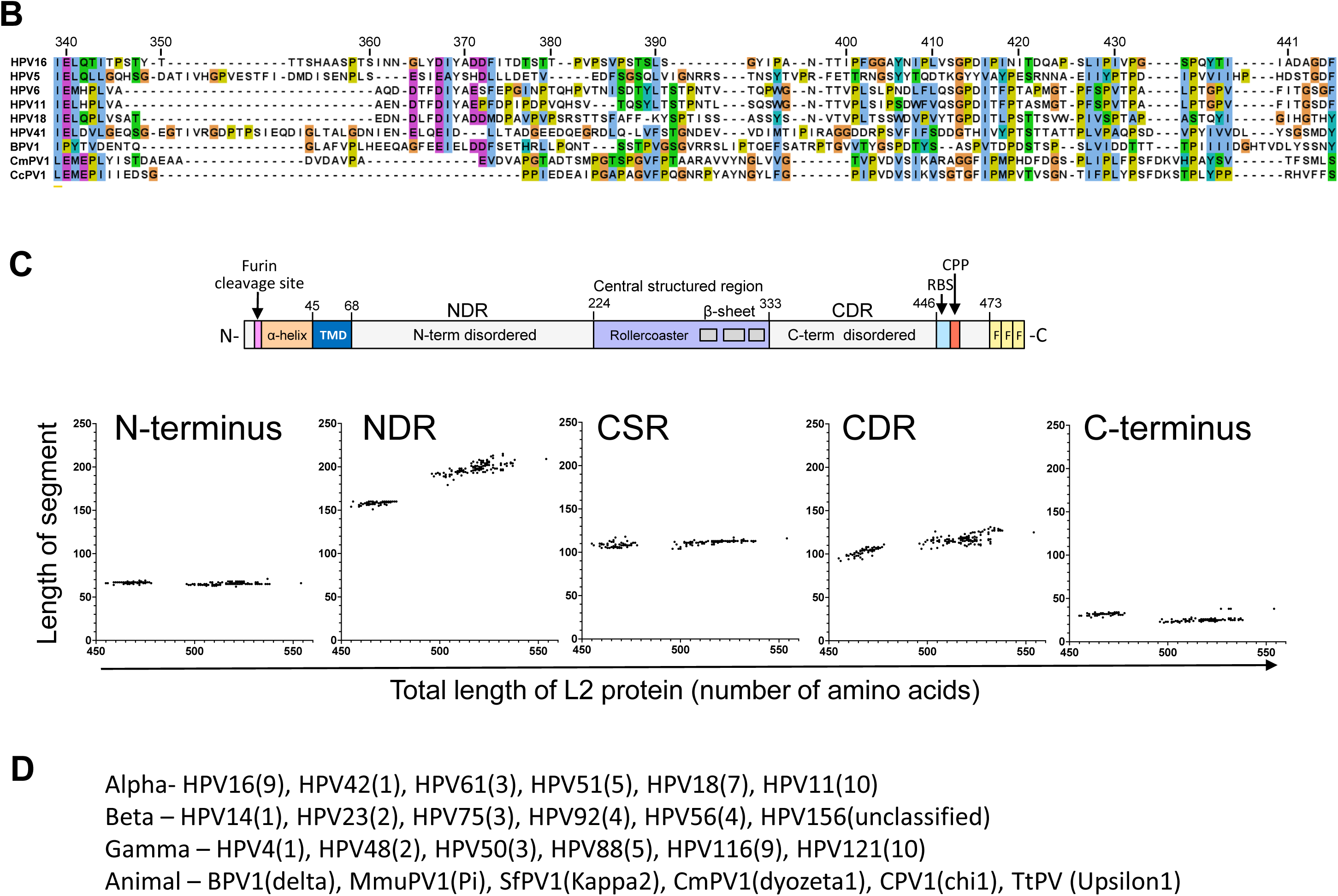
Phylogenetic map and sequence alignment of papillomaviruses. **A.** Papillomavirus phylogenetic tree based on an alignment of the DNA sequence coding for E1, E2, L1 and L2 proteins from 241 different papillomaviruses. AlphaFold2 was used to model L2 from the papillomavirus types indicated in red. Figure modified from (Van Doorslaer, 2013). **B.** Sequence alignment of HPV16 L2 (amino acid positions indicated at the top) with the corresponding segment of representative papillomaviruses, generated with Jalview (Waterhouse et al., 2009) **C.** Top, schematic diagram of the HPV16 L2 protein. Bottom. Graph showing the length of each segment in all sequenced HPV types, plotted vs. the total length of the L2 protein for each HPV type. Each spot represents a different HPV type. N-terminus, N-terminal segment segment through the TMD: NDR, N-terminal disordered region; CSR, central structured region; CDR, C-terminal disordered region; C-terminus, segment from position 446 to the C terminus of the L2 protein. **D.** List of papillomaviruses used to calculate L2 conservation in Figure 1C.

**Figure S2.**
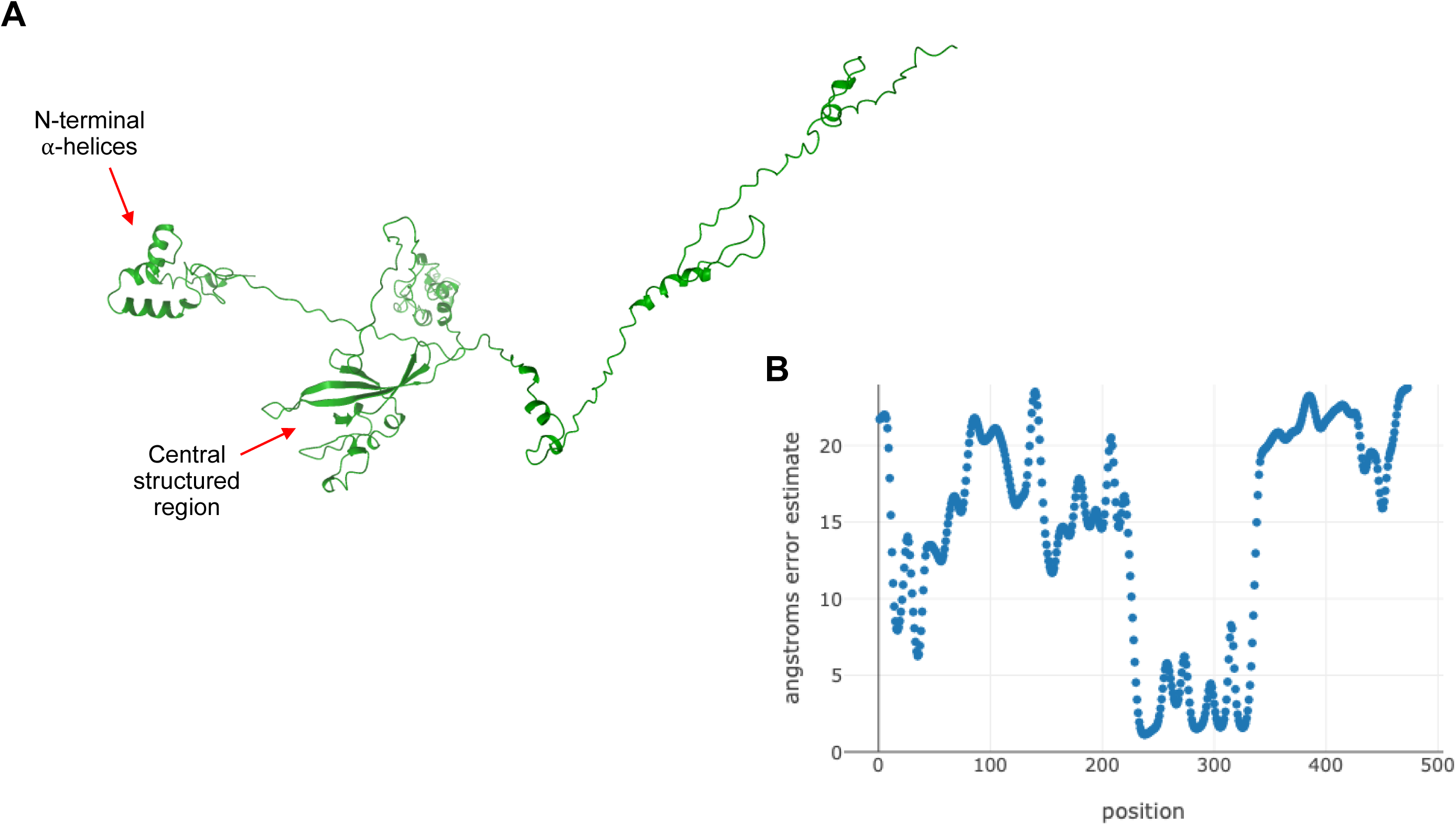
RoseTTAFold model of L2. **A.** RoseTTAFold predicted model for HPV16 L2. **B.** The Angstroms error estimate plot for the model shown in panel A. Lower values indicate higher in the prediction.

**Figure S3.**
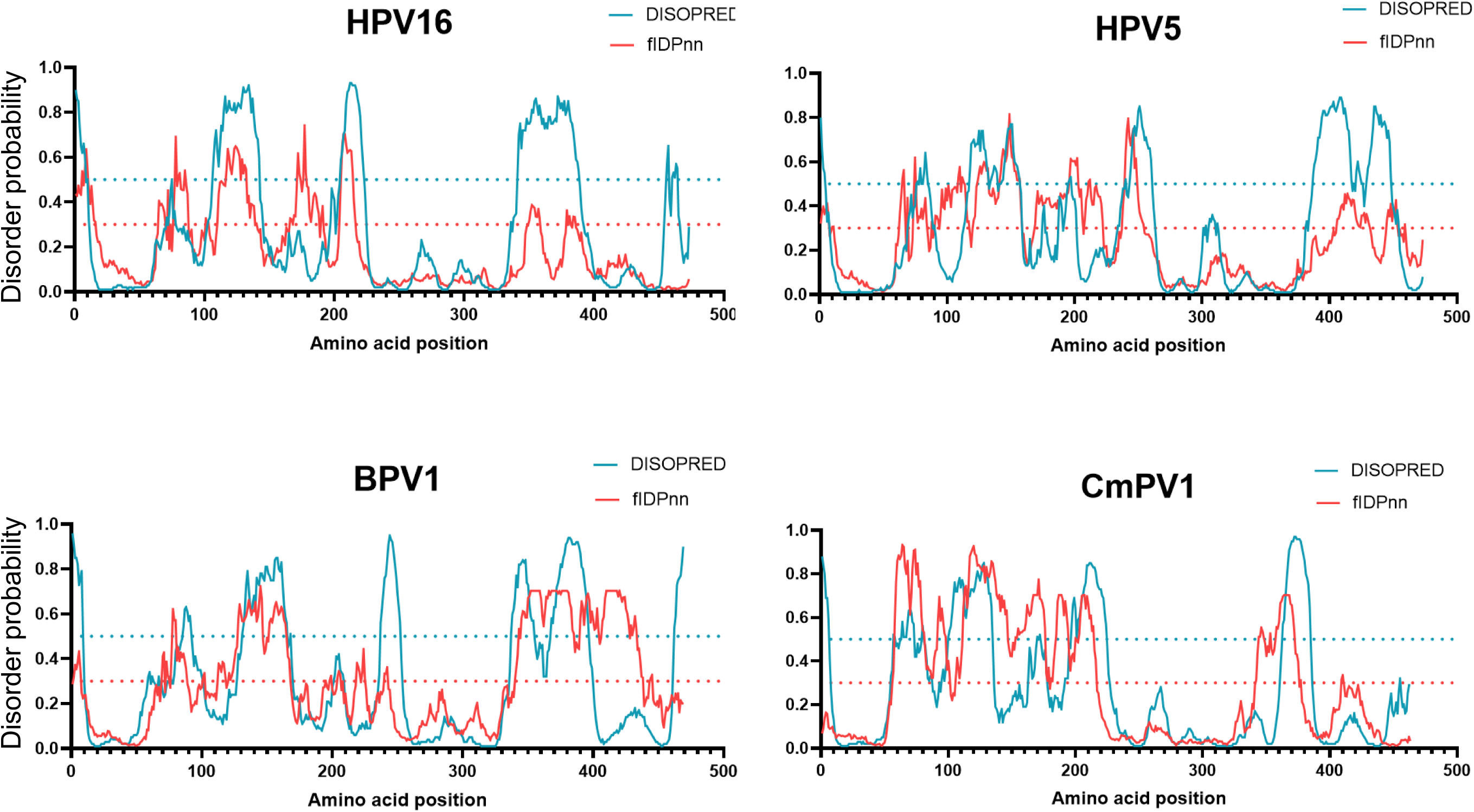
Predicted disordered region of various papillomavirus types. Intrinsic disorder profile of the L2 protein of the indicated papillomavirus types. Solid lines are DISOPRED (blue) and fIDPnn (red) prediction scores for L2. A higher value indicates higher probability of disorder. Horizontal dotted lines are the order/disorder threshold for DISOPRED (red) and fIDPnn (blue).

**Figure S4.**
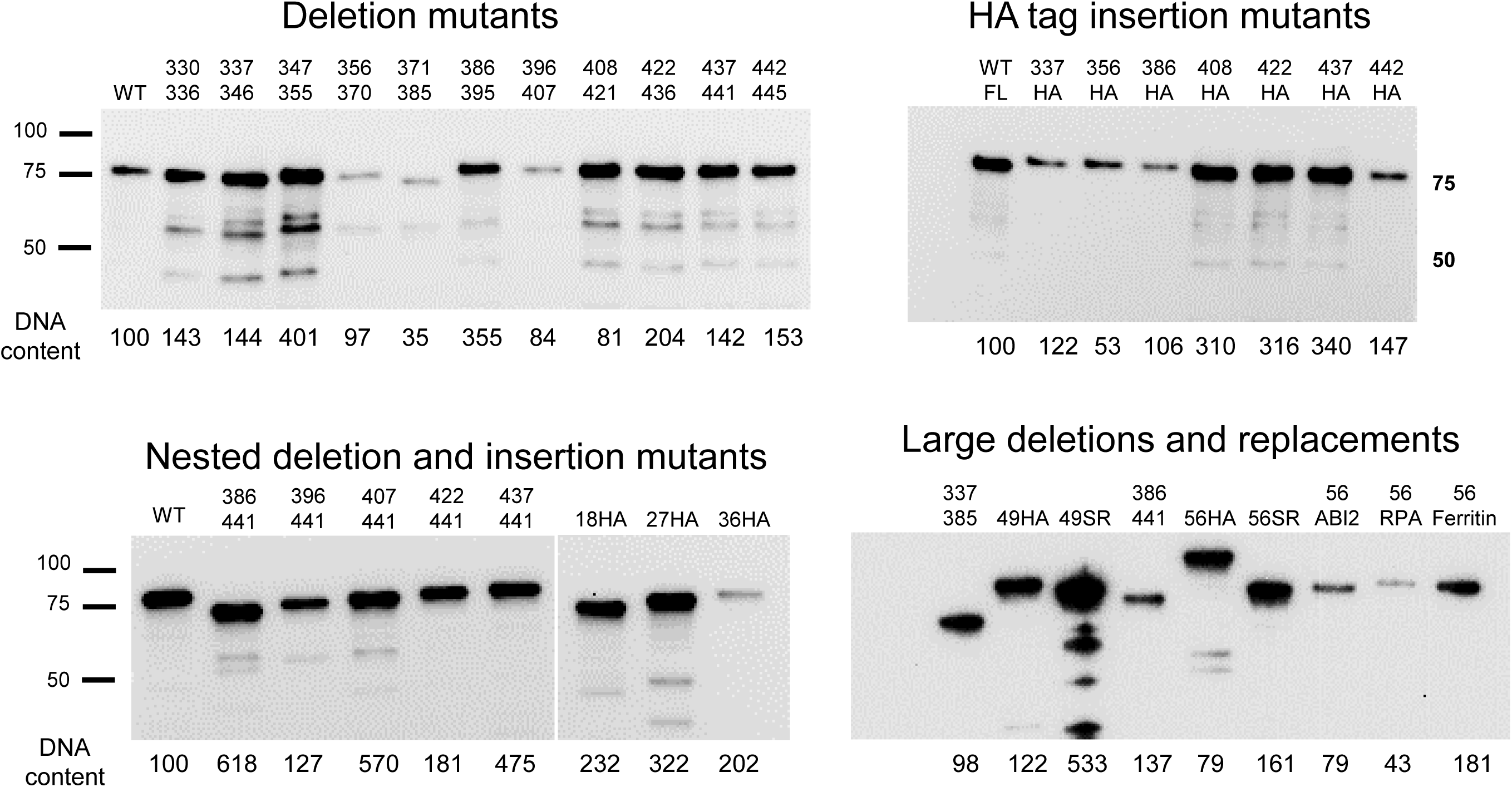
L2 and DNA content of pseudovirus stocks. Western blot analysis of representative wild-type and mutant HPV PsV purified by density gradient centrifugation. For each PsV, gradient fractions containing peak L1 levels as determined by SDS-polyacrylamide gel electrophoresis and Coomassie blue staining were pooled, and an equal volume of each pooled sample was subjected to electrophoresis. The images show western blots probed with anti-FLAG monoclonal antibody to detect L2. Reporter plasmid content of each PsV preparation was determined by quantitative polymerase chain reaction (qPCR) and expressed below each lane as percentage relative to the amount of DNA in the wild-type PsV preparation (set at 100%). For each experiment measuring infectivity, cells were infected with the same amount of encapsidated DNA that generated an MOI of 1 when wild-type PsV was tested.

**Figure S5.**
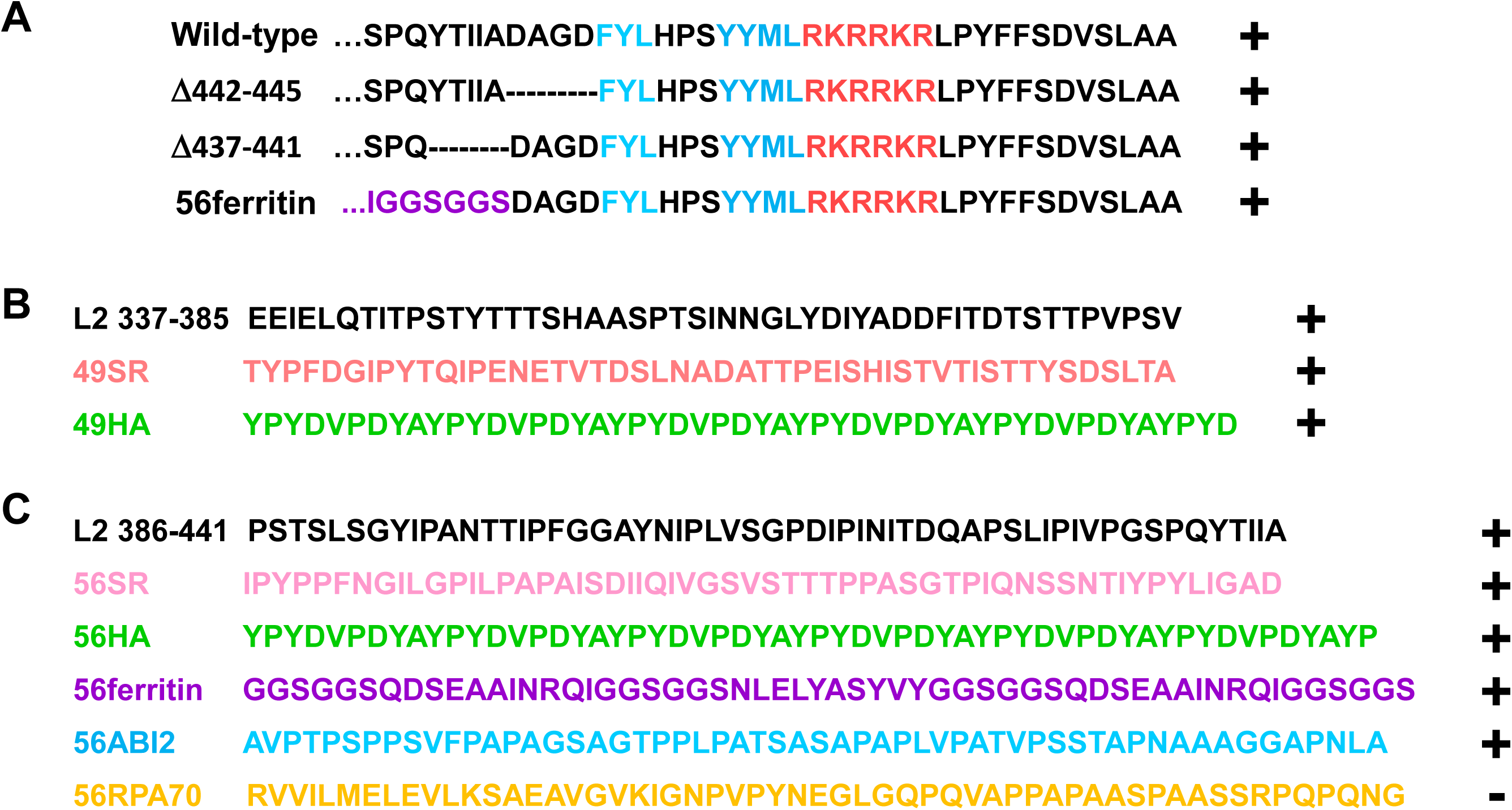
Amino acid sequence of wild-type and mutant HPV16 L2 segments. **A.** Sequence of amino acids from position 434 to the C terminus of the wild-type HPV16 L2 protein and the indicated infectious mutants. The FYL major retromer binding site and YYML minor retromer binding site are shown in blue and the cell-penetrating peptide is shown in orange. A portion of the segment replaced in 56ferritin is shown in purple. Dashes represent deleted amino acids. **B.** Amino acid sequence of wild-type HPV16 L2 amino acids 337-385 (top line) and its replacement mutants. **C.** As in panel B except amino acids 386-441 are shown. In panels B and C, the colors of the replacement mutants correspond to the colors used in Fig. 3A. In all panels, + and – indicate infectivity of PsV containing the indicated mutants.

**Figure S6.**
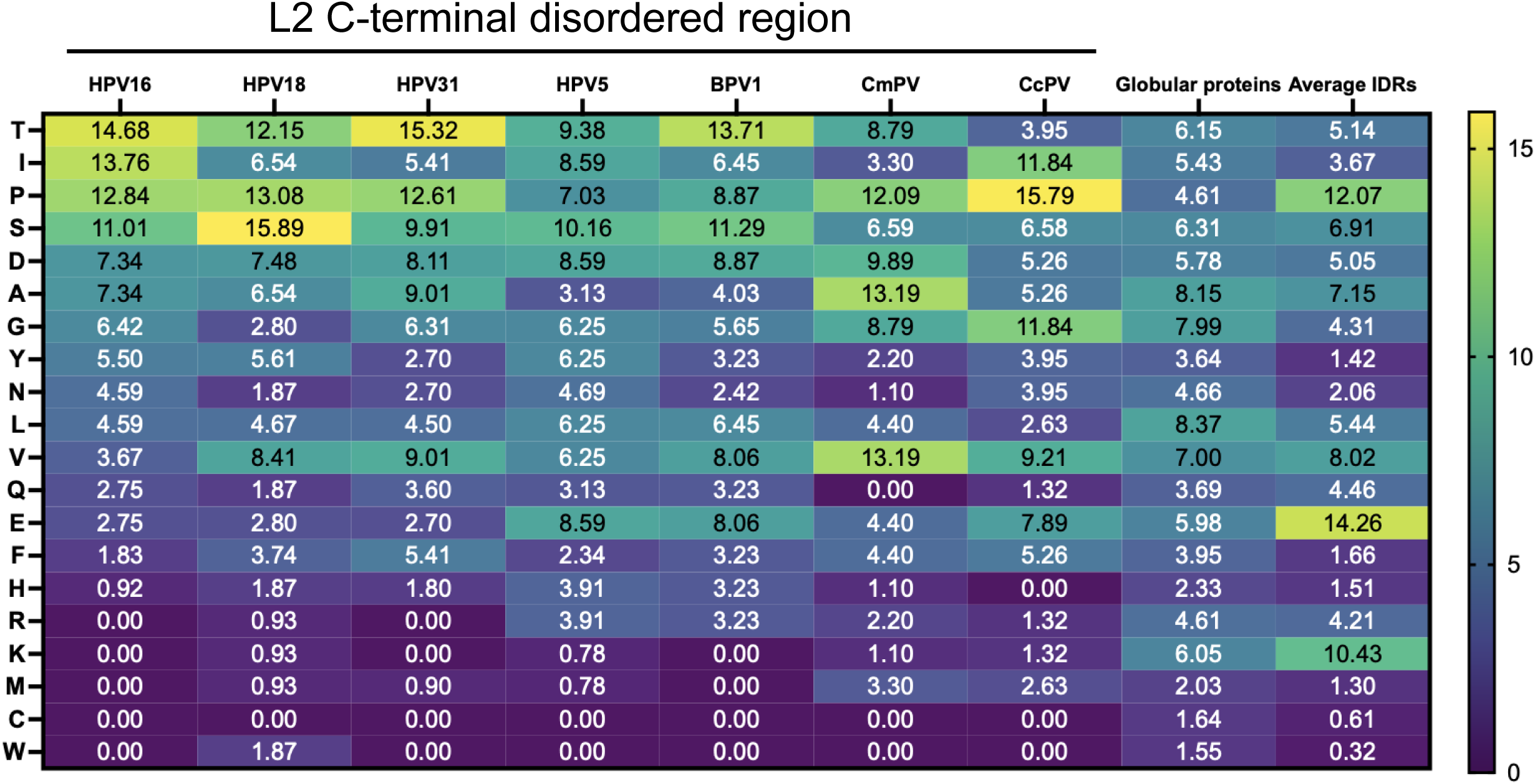
Amino acid composition of the L2 C-terminal disordered region of various papillomavirus types. C-terminal disordered region of HPV16 (positions 337-445) and the corresponding regions of other papillomavirus types. Amino acid composition of each region is plotted on a heatmap, shown in descending order based on abundance of each amino acid in the HPV16 L2 segment. Numbers show percent abundance of amino acid. For comparison, the average composition of large compilations of globular proteins and IDRs are shown (Tompa, 2002).

**Figure S7.**
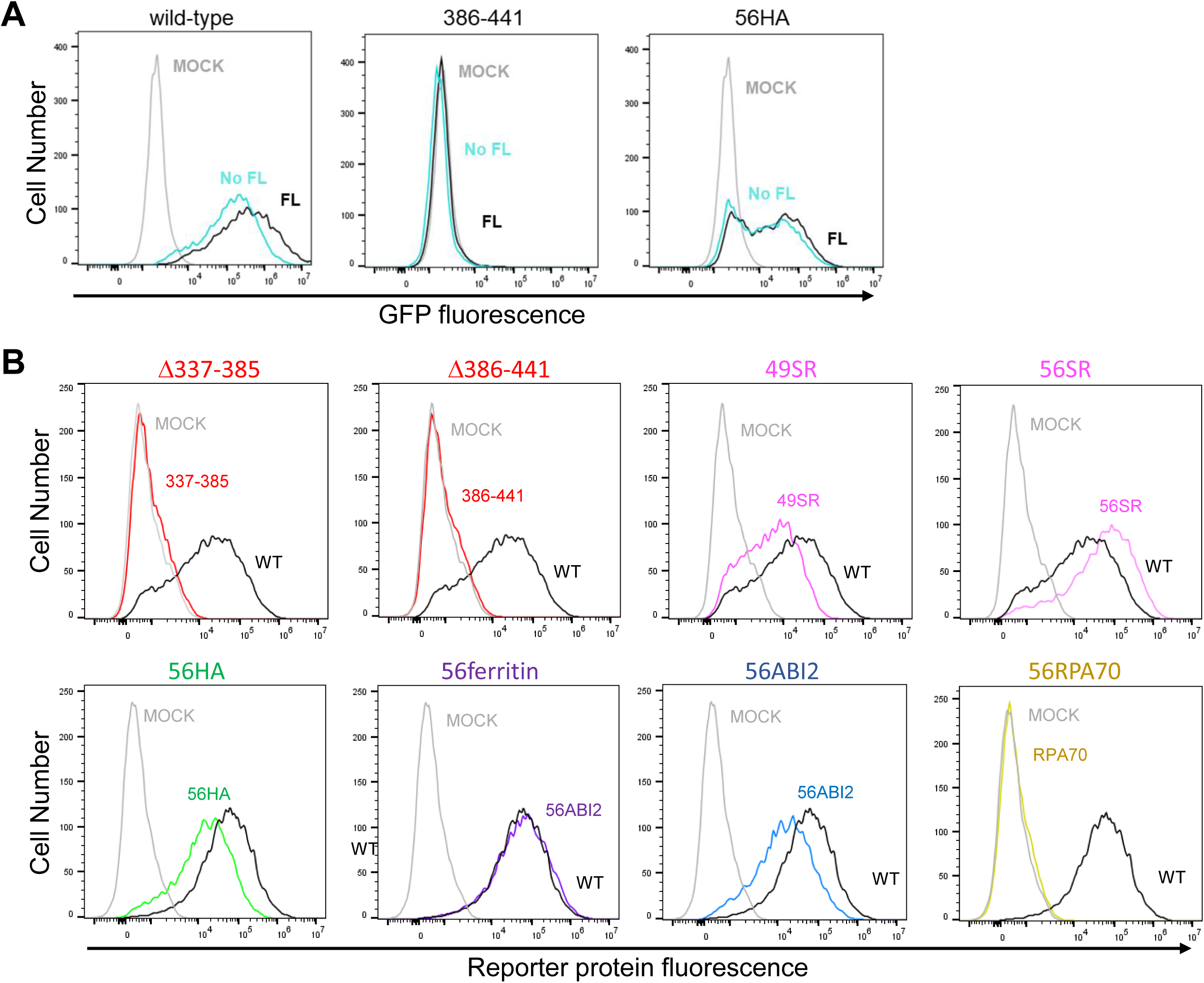
Infectivity of untagged PsV in HeLa cells and of FLAG-tagged PsV in HaCaT cells. **A.** HeLa S3 cells were mock-infected or infected in parallel at an MOI of 10 with PsV containing wild-type FLAG-tagged or untagged L2, or with tagged and untagged Δ386-441 and 56HA PsV stocks containing the same number of packaged reporter plasmids. Infectivity was assessed 48 hpi as described in the Fig 2. The graphs show histograms of a flow-cytometry experiment for the indicated PsV [mock infected, grey; FLAG-tagged (FL), black; untagged (no FL), light blue]. **B.** HaCaT cells were mock-infected or infected at an MOI of 100 with the indicated FLAG-tagged PsV. Infectivity was assessed 48 hpi as described in the Fig 2. The graphs show histograms of a flow-cytometry experiment for the indicated PsV (mock infected, grey; wild-type PsV, black; and L2 mutant, colored). In the top row, PsV expressed HcRed, and in the bottom row they expressed GFP. Similar results for each mutant were obtained in two experiments.

**Figure S8.**
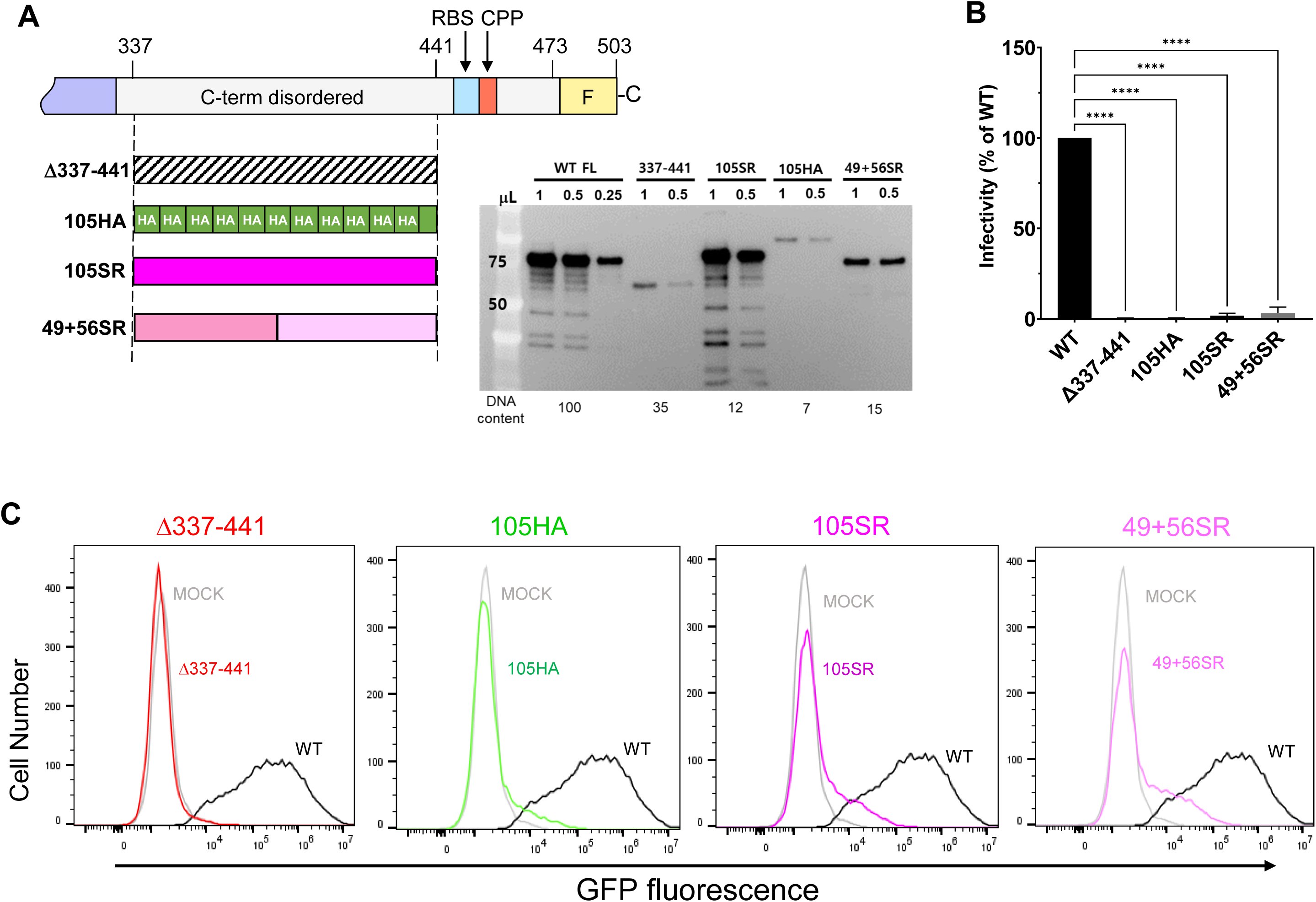
Replacement of entire C-terminal disordered region poorly rescues activity. A. Schematic diagram of the C terminus of wild-type and mutant HPV16 L2 proteins including Δ337-441 deletion mutant (hatched pattern), scrambled replacements (105SR and 49+56SR, pink), and tandem HA replacement (105 HA, green) mutants. Inset shows L2 western blot of different dilution of virus stocks and DNA quantitation of PsV preparations. **B.** Graph of the infectivity for wild-type and mutant PsV measured as in Figure 2. **C.** Flow cytometry histograms of infectivity measurements of mock-infected cells or cells infected with wild-type PsV or the indicated mutant. Note the shoulder of fluorescent cells in samples infected with the replacement mutants but not with the parental Δ337-441 deletion mutant.

**Figure S9.**
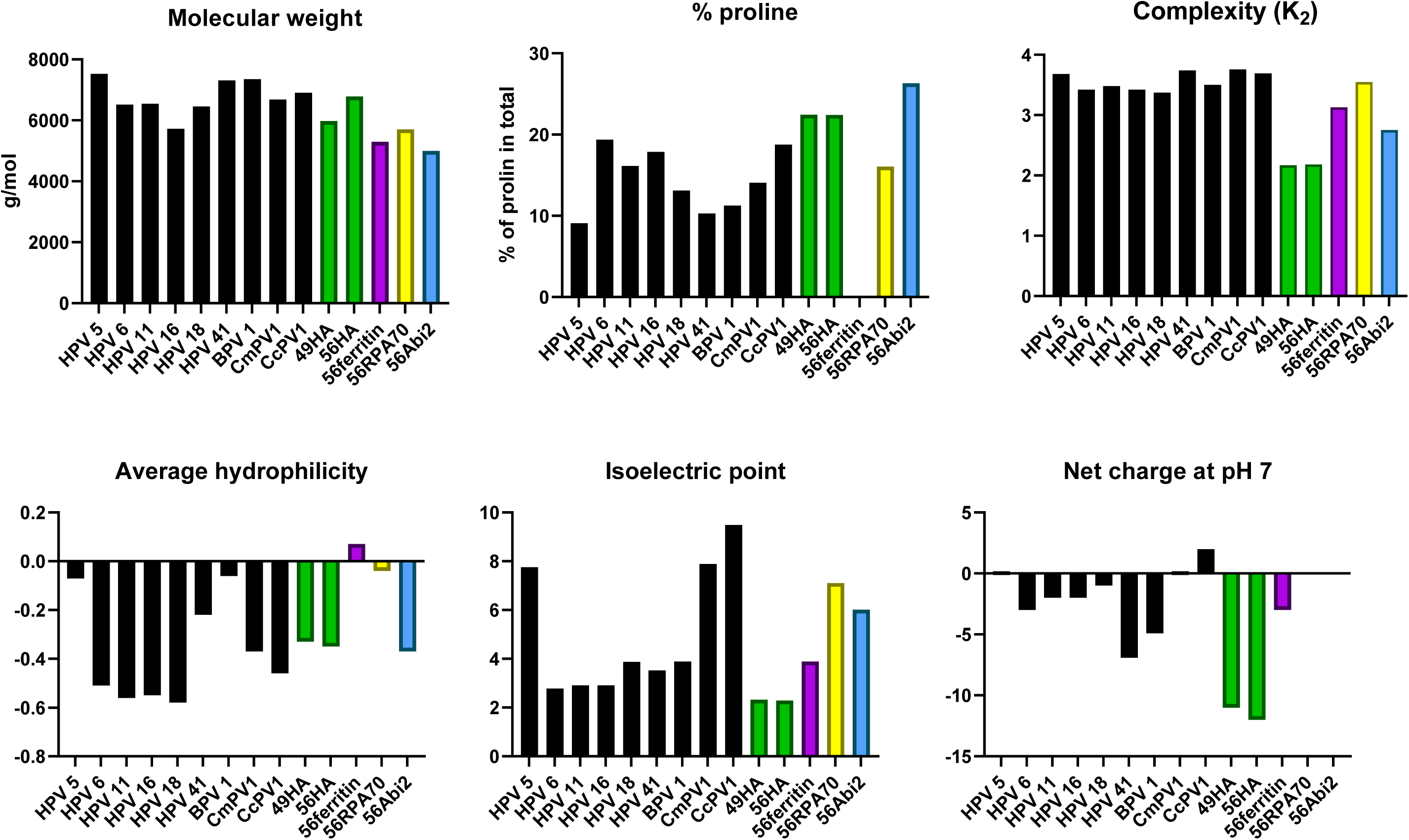
Biochemical properties of the C-terminal disordered region of L2 and the replacement mutants. Biochemical characteristics were calculated and graphed for the C-terminal disordered region of HPV16 L2 (amino acid positions 386-441) and of the corresponding regions in various papillomaviruses based on sequence alignment. The graphs also show the results for 49HA, 56HA, 56Abl2, and 56ferritin-helix, which functionally replaced the wild-type L2 segment, and for 56RPA70 (yellow bar), which did not. Sequence complexity scores (K_2_ values) were calculated as described by Wootton and Federhen (Wootton & Federhen, 1996). Average hydrophilicity score was calculated as described by Hoop and Woods (Hopp & Woods, 1981).

**Figure S10.**
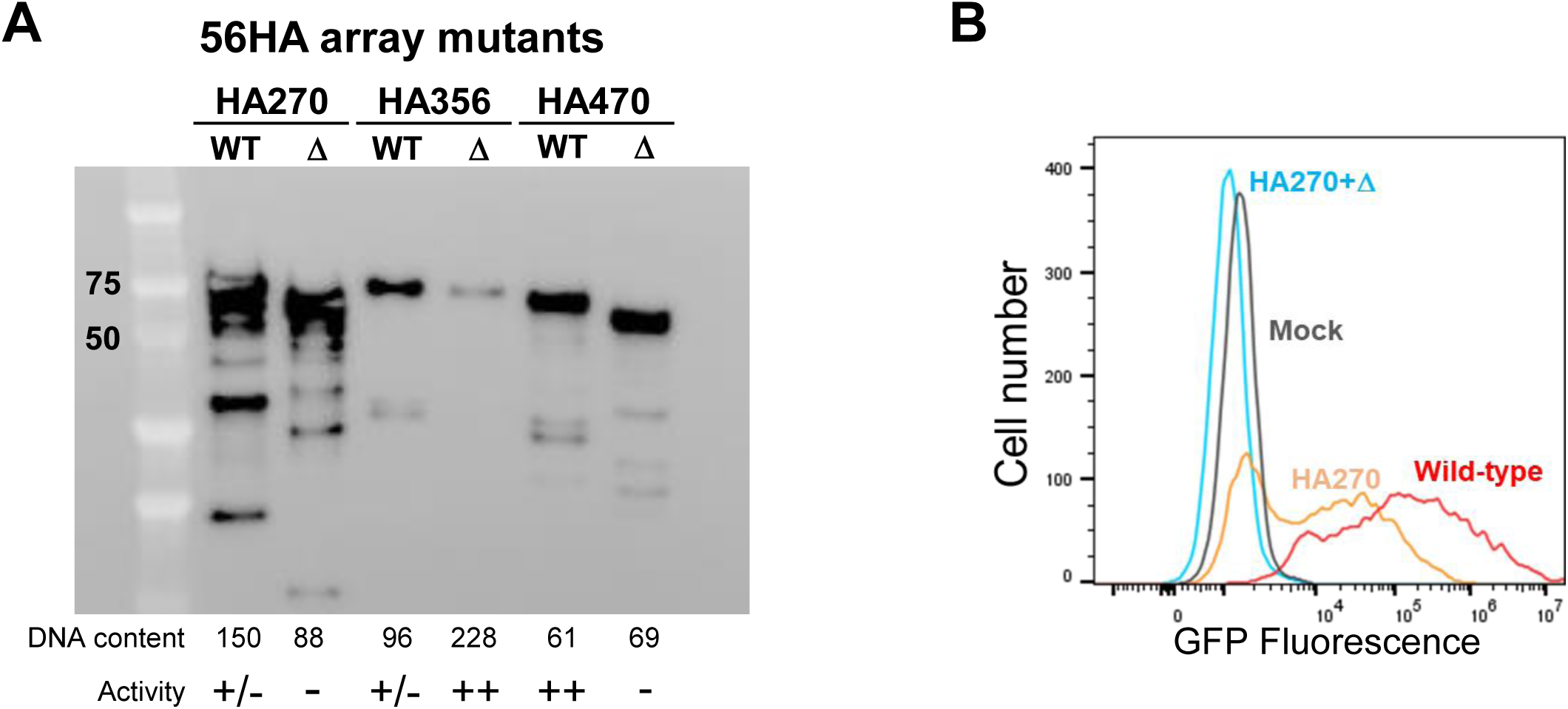
HA array at position 270 does not rescue the infectivity defect of Δ386-441. **A.** Western blot of L2 in purified PsV as in Figure S4. The relative infectivity of PsV containing the indicated L2 protein is indicated below the panel. **B.** Flow cytometry histograms of HeLa S3 cells 48 hpi at a MOI of 100 with wild-type PsV and PsV containing the 56-residue HA tag array at position 270 in the presence and absence of Δ386-441.

**Figure S11.**
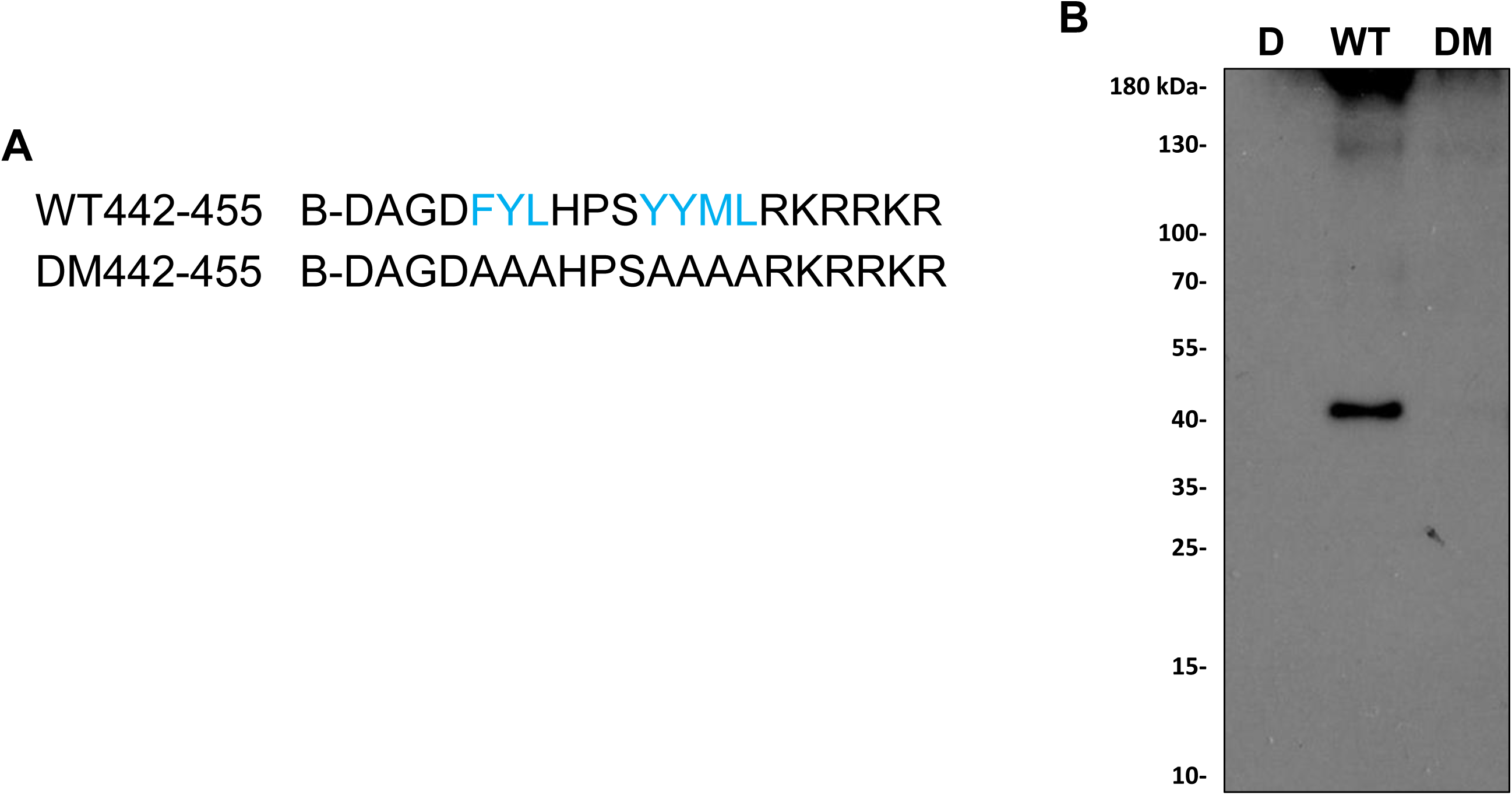
Binding of retromer to L2 deletion mutant peptide. **A.** Sequences of the wild-type and mutant L2 peptide (amino acids 442 to 461 of HPV16 L2) used in this study. B indicates N-terminal biotin. The retromer binding site is shown in blue. **B.** Western blot of peptide pull-down experiment. Wild-type and mutant biotinylated peptides were added to lysates of uninfected HeLa S3 cells. After incubation, peptides and associated proteins were pulled-down with magnetic streptavidin beads. After SDS-PAGE, retromer associated with peptide was detected by immunoblotting for VPS26. (D, DMSO control; WT, wild-type peptide; DM, retromer binding site mutant peptide.

**Figure S12.**
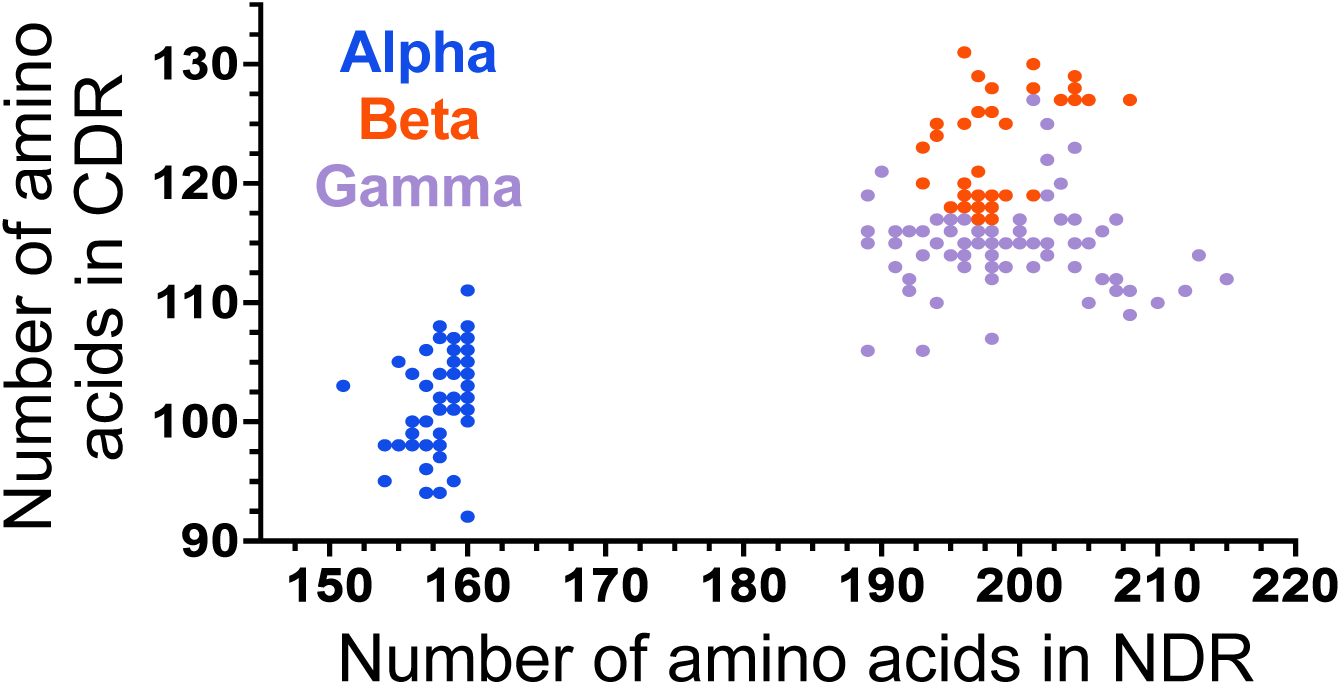
Comparison of the length of the disordered regions in L2 from different HPV types. Scatter plot showing the correlation between the number of amino acids in the predicted CDR vs. the number of amino acids in the predicted NDR for all sequenced HPV. Each spot represents a different HPV type, colored according to their phylogenetic clade.

## Materials and Methods

### Cells

HeLa S3 cervical carcinoma cells were purchased from American Type Culture Collection (ATCC). HaCaT cells, a spontaneously immortalized but non-tumorigenic adult human skin keratinocyte cell line that retains epidermal differentiation potential (Boukamp et al., 1988) were purchased from AddexBio Technologies. 293TT cells were generated by introducing SV40 Large T antigen cDNA into female HEK293T cells to increase Large T antigen expression and were obtained from Christopher Buck (National Institutes of Health). All cells were cultured at 37°C in with Dulbecco’s modified Eagle’s medium (DMEM, Gibco) with HEPES and L-glutamine, supplemented with 10% fetal bovine serum (FBS) and 100 units/mL penicillin streptomycin, in 5% CO_2_.

### Plasmids

Packaging plasmids designated p16sheLL, expressing HPV16 L1 and L2, were used to produce wild-type or mutant PsV (Buck et al., 2005).To facilitate cloning, Q5 site-directed mutagenesis protocol was used to introduce five restriction enzyme cleavage sites (BstXI, PflFI, BmtI, BbvCI, and XhoI) within the L2 coding region and downstream of the L2 gene. None of the mutations affected the amino acid sequence of L2 except BstXI (Serine 219 to Threonine), which did not affect PsV infectivity. The L2 mutants were constructed in p16sheLL by cloning synthetic DNA (gBlocks, IDT) containing the mutations between pairs of these cleavage sites. The entire L2 gene in each mutant was sequenced to verify the presence of the desired mutations and the absence of inadvertently introduced mutations. For mutants displaying a severe infection defect, the sequence of the wild-type L1 gene was also confirmed. Details of restriction site introduction and sequences of mutations are available upon request from the authors.

### Pseudovirus production and characterization

To produce PsVs, 5 x 10^6^ 293TT cells were seeded per plate in 15 cm dishes and cultured for 24 h before transfection. Polyethyleneimine was then used to co-transfect the cells with reporter plasmid [pCINeo-GFP (obtained from Christopher Buck, NIH) or pCAG-HcRed (Addgene #11152)] and p16sheLL expressing HPV16 L1 and wild-type or mutant L2. Packaged PsVs were purified by density gradient centrifugation in Optiprep and gradient fractions containing the most L1 protein were pooled (Buck et al., 2005).The presence of viral proteins in PsV stocks was assessed by SDS-polyacrylamide gel electrophoresis and Coomassie brilliant blue staining or immunostaining for L1 and L2 levels, respectively. Encapsidated reporter plasmids in PsV stocks were quantified by qPCR as described with modifications (Popa et al., 2015). Briefly, purified PsVs were treated with DNase I (PROMEGA) to remove unencapsidated DNA associated with capsids, followed by heat-inactivation of DNase I. Samples were then treated with 1.15 μg/mL proteinase K in the presence of 40 μM DTT, 40 μM EDTA and 50 ng/μL salmon sperm shredded DNA (sssDNA, ROCKLAND). The reporter plasmid was isolated using DNeasy Blood and Tissue Kit (QIAGEN), and the isolated plasmids were linearized with BglII and the copy number of encapsidated reporter plasmid was determined by qPCR using appropriate primers for the reporter gene in comparison to a standard curve.

### Measuring Infectivity

To measure infectivity of PsV stocks, 10^5^ HeLa S3 cells or HaCaT cells per well in 12 well plates were incubated with wild-type PsV at infectious multiplicity of infection (MOI) of 10 (HeLa cells) or 100 (HaCaT cells), as determined by flow cytometry for reporter protein expression in HeLa cells. The number of packaged reporter plasmids required to achieve this MOI for wild-type PsV was quantified by qPCR, and mutant PsV stocks containing the same number of encapsidated reporter plasmids were used to infect cells in parallel with wild-type. Approximately 48 hpi, cells were analyzed by flow cytometry on Cytoflex LX to determine the mean fluorescence intensity (MFI). Infectivity of mutants is expressed as the MFI induced by the mutant as a percentage of MFI induced by wild-type (set at 100%).

### Proximity ligation assay (PLA)

For PLA, 0.4-0.5 x 10^5^ HeLa S3 cells per well were plated in 24-well plates containing glass coverslips 24 h prior to infection. Cells were mock-infected or infected at MOI of ∼200 with wild-type PsV containing a reporter plasmid expressing HcRed (to eliminate possible interference of the fluorescent signal generated by GFP and PLA), as determined by flow cytometry for HcRed expression. The amount of L1 required to achieve this MOI for wild-type PsV was quantified by SDS-PAGE and Coomassie brilliant blue staining, and mutant PsV stocks containing the same amount of L1 in mutant PsV stocks were used to infect cells in parallel with wild-type. At indicated times post-infection, cells were fixed with 4% paraformaldehyde (Electron Microscopy Sciences) at room temperature for 12 min, permeabilized with 1% Saponin (Sigma-Aldrich) at room temperature (RT) for 35-40 min, and blocked using DMEM containing 10% FBS at RT for 1-1.5 h. Cells were then incubated overnight at 4°C with a pair of antibodies – mouse antibodies recognizing L1 (BD Biosciences, 554171, 1:1,000 when used with anti-TGN46, 1:200 when used with other antibodies) and a rabbit antibody recognizing a cellular marker protein (anti-TGN46, Abcam, ab50595, 1:600; anti-EEA1, Cell Signaling Technology, 2411, 1:75; anti-VPS35, Abcam, ab157220, 1:200). PLA was performed with Duolink reagents (Sigma Aldrich) according to the manufacturer’s instructions as described (Lipovsky et al., 2013). Briefly, cells were incubated in a humidified chamber at 37°C with a pair of PLA antibody (mouse and rabbit) probes for 75 min, with ligation mixture for 45 min, and then with amplification mixture for 3 h, followed by series of washes. Nuclei are stained with 4,6-diamidino-2-phenylindole (DAPI). Fluorescence from PLA in each cell was imaged using the Zeiss LSM980 confocal microscope. Images were processed using a Zeiss Zen software version 3.1 and quantified using Image J Fiji version 2.3.0/1.53f.

### Peptide binding experiments

N-terminal biotinylated peptides were synthesized by Genscript at ≥95% purity and dissolved in 100% DMSO to a stock concentration of 10 µg/µL. Lysates of uninfected HeLa S3 cells were prepared by plating 1.5-1.75 x 10^6^ cells in 6 cm dishes and incubating at 37°C for 12 h. Cells were trypsinized and collected before being lysed in 165 µL HEPES buffer (20 mM HEPES pH 8.0, 50 mM NaCl, 5 mM MgCl_2_, 1 mM DTT, and 1% Triton X-100) supplemented with 1X HALT protease and phosphatase inhibitor cocktail (ThermoFisher) for 45 min on ice. Lysates were clarified at 14,000 x g for 15 min at 4°C. The supernatant was collected and diluted with HEPES lysis buffer to 500 µL total to create the input sample. Input lysate was divided into separate tubes and incubated with DMSO or 10 µg of each biotinylated peptide for 2 h at 4°C. For pull down, 50 µL of Pierce™ Streptavidin Magnetic Beads (ThermoFisher) were equilibrated with HEPES lysis buffer and added to each tube. Beads were incubated for 1 h at 4°C and then washed 4X with HEPES lysis buffer. 2X Laemmli buffer with DTT was added and samples were boiled to elute proteins. Samples were analyzed by SDS-PAGE and immunoblotted with VPS26 antibody (Abcam #ab23892).

### Selective permeabilization

To measure L2 protrusion following selective permeabilization, 0.5 x 10^5^ HeLa S3 cells were seeded per chamber in 8-chamber glass slides (NUNC) 24h prior to infection. Cells were then infected with PsVs containing FLAG-tagged wild-type or mutant L2 at MOI of 100 (normalized as described in the Measuring Infectivity section). At 8 hpi, cells were washed with phosphate-buffered saline (PBS) at pH 10.7, fixed for 10 min at RT with 4% paraformaldehyde and washed with PBS. Cells were permeabilized with 1% (W/V) saponin (Sigma-Aldrich) in PBS for 40 min at RT or selectively permeabilized with freshly dissolved digitonin (2μg/ml) in PBS for 10 min at RT, washed with PBS, and blocked with DMEM supplemented with 10% FBS and 5% normal donkey serum for 1 hour at RT. Cells were then incubated with 1:1000 anti-L1 mouse antibody (BD, 554171) or 1:250 anti-FLAG rabbit antibody (Cell Signaling Technology, D6W5B) at RT overnight. Cells were then incubated at RT for 2 hours with 1:1000 Alexa Fluor (AF) 647 donkey anti-mouse and 1:200 of AF488 donkey anti-rabbit fluorescence conjugated secondary antibodies (Life Technologies). The cover slip was mounted in mounting solution with DAPI, and images were taken using Zeiss LSM 980 confocal microscope. Images were processed using a Zeiss Zen software version 3.1 and quantified using Image J Fiji version 2.3.0/1.53t software. Similar results were obtained in three independent experiments for each pair of antibodies.

### Structure and disorder predictions

Structures of the L2 protein were predicted using AlphaFold2 (Jumper et al., 2021) on the Google Colab notebook with default settings or with RoseTTAFold on the Robetta server (Baek et al., 2021). The ColabFold Google Colab notebook was also used to generate multiple ranked structural models of the L2 protein with AlphaFold2 (Mirdita et al., 2022). Similar results were observed for all ranks. Predicted structures were visualized using ChimeraX and Mol*(Pettersen et al., 2021) (Sehnal et al., 2021). pLDDT values were obtained from the AlphaFold2 prediction, and the sequence conservation of L2 was calculated from an alignment of 18 HPV types and six animal papillomaviruses by Jalview (Waterhouse et al., 2009) and averaged over a five amino acid window. Disorder probability was predicted using DISOPRED3 accessed through the PSIPRED server (McGuffin et al., 2000) and using flDPnn (Hu et al., 2021).

AlphaFold Multimer was run by using XSEDE supercomputing resources (Cianfrocco et al.; Evans et al., 2022). The modeling includes VPS26 (aa 8-301, ID:O75436-1), VPS35 (aa 12-363, ID: NP_060676.2), SNX3 (aa 1-160, ID: O60493.3), and HPV16 L2 (aa 444-456, ID: P03107). The model does not include the C-terminal 13 helices of VPS35 or VPS29 because this part of the complex is not involved in cargo recognition. The inner and outer leaflets of the membrane are shown as cylindrical masks derived from the cryo-ET map (EMD-12221) of the retromer:SNX3 membrane coat (Leneva et al., 2021).

### Biochemical property calculation

Biochemical properties of protein segments were calculated using BACHEM peptide calculator (https://www.bachem.com/knowledge-center/peptide-calculator/). Sequence complexity scores (K_2_ values) were calculated as described by Wootton and Federhen (Wootton & Federhen, 1996). Average hydrophilicity score was calculated as described by (Hopp & Woods, 1981).

